# Enteric sensory neurons for nutrient detection and gut motility

**DOI:** 10.64898/2026.03.04.709708

**Authors:** Kai Li, Junhui Mou, Xiaohan Sun, Yuanyuan Chen, Lin Fu, Zhuang Wang, Yaru Wei, Meirong Wang, Peng Guo, Xiaolin Lin, Liang Wang, Shumin Duan, Stephen D. Liberles, Jinfei D. Ni

## Abstract

The enteric nervous system (ENS) orchestrates gastrointestinal reflexes and brain-gut communication via molecularly diverse neurons. Among these, intrinsic primary afferent neurons (IPANs) are essential for detecting luminal nutrients and irritants, yet their molecular identities, sensory properties, and functions remain poorly resolved. Here, we establish a segment-resolved single-cell atlas of the murine ENS, including a comprehensive characterization of the gastric ENS. This resource defines a refined taxonomy of enteric neurons and glia and enabled the development of a genetic toolkit for molecularly defined IPANs. Using chemogenetics and calcium imaging, we discovered that myenteric neurons detect a wide range of nutrients, irritants, and cytokines. Nutrient detection depends on a functional connection between chemosensory epithelial cells and enteric neurons mediated by 5-HT—HTR3 axis. Through optogenetic analysis, we demonstrated segment-specific regulation of gut motility by different IPANs. Our work establishes a genetic and physiological framework for enteric-specific sensory mechanisms.

## Introduction

The enteric nervous system (ENS) participates in most aspects of gastrointestinal physiology including gastric empty, food digestion and absorption, defecation and gut immunity^1–4^. Together with the gut-innervating peripheral sensory neurons and autonomic motor neurons, they form the basis of the gut-brain bidirectional neural connection^5^. Dysfunction of the ENS is implicated in numerous gastrointestinal disorders, including Hirschsprung’s disease, gastroparesis, chronic constipation, and irritable bowel syndrome, highlighting its critical role in maintaining gut homeostasis^2,6^.

In mammals, the enteric nervous system is composed of interconnected neural network distributed in two enteric plexuses: myenteric plexus sandwiched between longitudinal and circular smooth muscle layers and submucosa plexus beneath the gut mucosa^1,7^. Over the past 50 years, studies have classified the mammalian ENS using immunohistochemical, neurophysiological, and pharmacological approaches. These efforts established a model in which the ENS network consists of intrinsic primary afferent neurons (or IPANs), interneurons, motor neurons, and associated glia, each with distinct neurochemical, morphological, and physiological properties^2,8,9^. Recent advances in single-cell transcriptomics, combined with genetic and viral tools, have further elucidated ENS diversity, development, and function^10–12^. However, existing studies remain limited and incomplete—for example, the gastric ENS has yet to be thoroughly characterized at the molecular level. To fully harness the potential of neuronal genetics in model organisms, a tailored pipeline is needed for comprehensive ENS profiling.

The gastrointestinal tract continuously monitors a vast array of chemical and mechanical stimuli to orchestrate essential physiological processes and behavioral responses^3,13–16^. This sensory capability is mediated by specialized neurons originating from three distinct anatomical sources: (1) vagal sensory neurons residing in the nodose ganglia^17–20^, (2) spinal sensory neurons located in the dorsal root ganglia^21–24^, and (3) IPANs embedded within the ENS. Recent advances in genetic targeting strategies have revolutionized our understanding of peripheral sensory pathways, enabling comprehensive characterization of vagal and spinal sensory neurons with unprecedented resolution^20,25–29^. In contrast, while the existence of IPANs as the intrinsic sensory arm of the ENS has been demonstrated since the 1990s, our understandings of their fundamental properties, including their sensory response properties, neurochemical coding, anatomical organization, morphological diversity, and physiological functions, remain limited^8,30–34^. Genetic tools to access IPANs are critical to interrogate these fundamental questions.

In this study, we developed an efficient pipeline of single-cell transcriptomic analysis for a large majority of the gastrointestinal tract (stomach, small intestine, and colon), establishing a comprehensive taxonomy of both enteric neurons and enteric glia. Sweeping through a collection of Cre or FlpO reporter lines based on genetic markers identified from our analysis, we established a genetic toolkit for interrogating the anatomy, morphology and sensory response properties of putative IPANs. Combining chemogenetics and calcium imaging, we discovered that myenteric neurons could detect a wide range of nutrients, irritants, and cytokines. Nutrient detection depends on a functional coupling between chemosensory epithelial cells and enteric neurons mediated by 5-HT. Our optogenetic assisted functional analysis revealed differential regulation of gut motility by enteric subpopulations. Our work established a molecular and physiological framework to investigate sensory-motor responses mediated enteric sensory neurons.

## Results

### A comprehensive cell atlas of the mice enteric nervous system revealed by single-cell RNA sequencing

To establish a comprehensive cellular taxonomy across gastrointestinal segments, we developed a pipeline involving for segment-specific single-cell transcriptomics of the murine enteric nervous system (ENS). Our strategy employed three complementary transgenic approaches (**Figures 1a-c, Extended Data Figure 1a-d**) to overcome tissue-specific technical constraints (See method part for rational and details). In total, we analyzed single-cell transcriptomes from 83555 cells of the murine ENS (**Extended Data Figure 1e**), including 21810 cells from the stomach myenteric plexus, 20123 cells from the small intestine myenteric plexus, 4987 cells from the small intestine submucosal plexus, and 36635 cells from the colon myenteric plexus. Clustering analysis was performed independently for each tissue, and enteric neurons or glia were assigned based on established markers (**Extended Data Figure 1f-i**): *Snap25* and *Elavl4* for enteric neurons, and *S100b* and *Gfap* for enteric glia.

**Figure 1.**
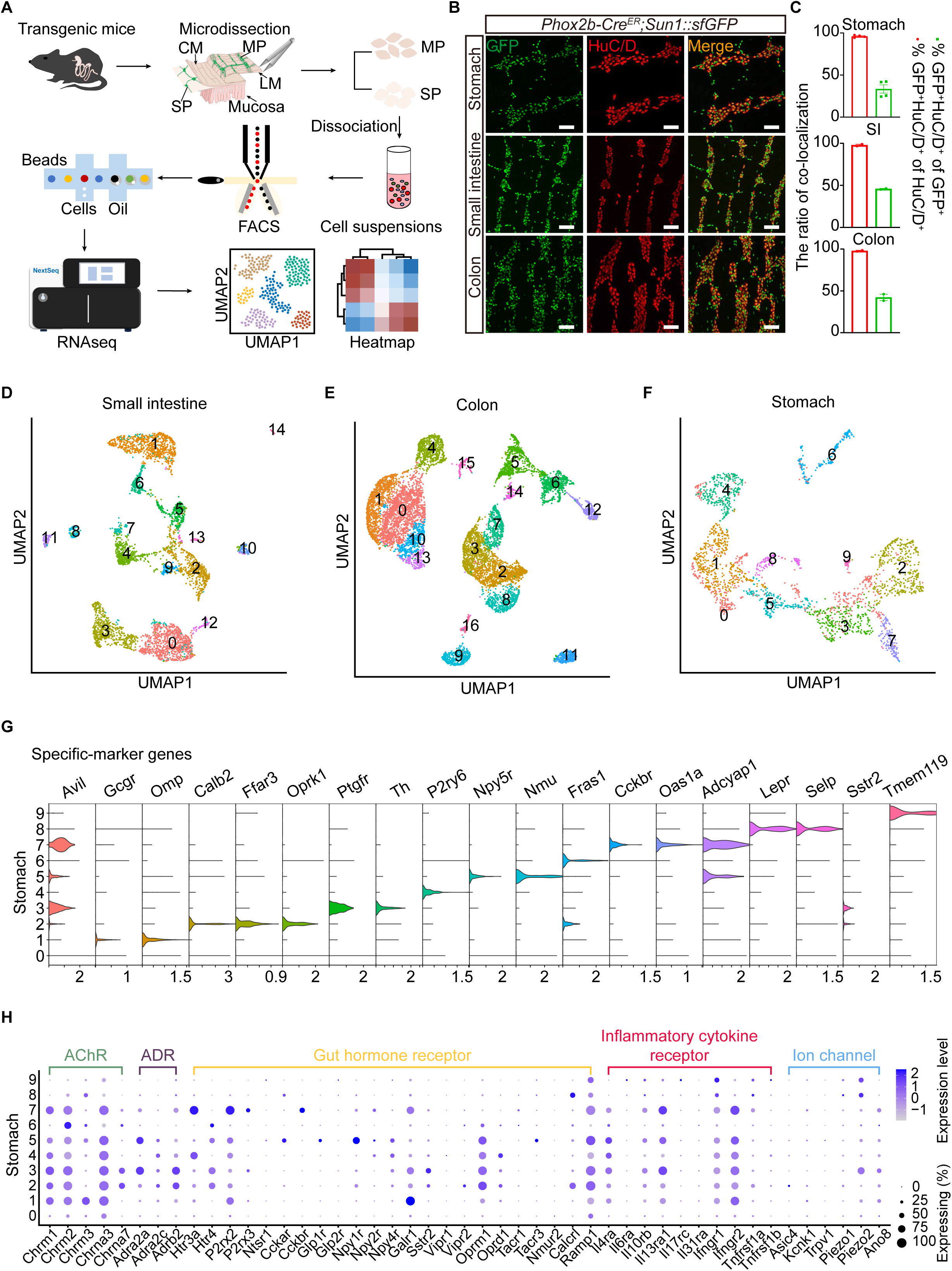
A scRNA-seq pipeline for segment specific profiling of the mouse ENS. **(a)** Schematic diagram of the workflow for the scRNA-seq pipeline. **(b-c)** Staining and quantification of cells in the myenteric plexus labeled by *Phox2b-Cre^ER^* driven nuclei-GFP. HuC/D is used to label enteric neurons. Data are presented as mean ± SEM. Scale bar, 100 µm. (n = 4 for stomach, n = 2 for small intestine and colon). **(d)** Uniform manifold approximation and projection (UMAP) of small intestine myenteric neurons, color-coded to represent 15 distinct clusters. **(e)** UMAP of colon myenteric neurons, color-coded to represent 17 distinct clusters. **(f)** UMAP of stomach myenteric neurons, color-coded to represent 10 distinct clusters. **(g)** Violin plot showing marker genes for 10 subsets of stomach myenteric neurons. **(h)** Dot plots showing differentially expressed genes categorized as acetylcholine receptor (AChRs), adrenergic receptor (ADRs), gut hormone receptors, inflammatory cytokine receptors, and ion channels of enteric neurons in stomach. The color scale represents the gene expression, and dot size represents the percentage of cells.

We integrated single-cell RNA sequencing (scRNA-seq) datasets according to tissue of origin and computationally isolated enteric neurons and glia for separate analysis. Re-clustering analysis revealed 12 enteric glia subtypes (**Extended Data Figure 2**). Subsequent clustering of enteric neurons revealed 15 transcriptionally distinct myenteric neuron subtypes in the small intestine and 17 in the colon (**Figures 1d–e**). Our analysis recapitulated major subtypes defined in prior studies of myenteric neurons (comparative analysis: **Extended Data Figure 3a–b**). Based on marker genes used to functionally categorize myenteric neurons^10–12^, our small intestine dataset composed of 8 classes of motor neurons (4 excitatory, 4 inhibitory), 2 classes of interneurons, 4 classes of IPANs, and an unknown class (**Extended Data Figure 3c**). This refined previous categorization based on juvenile myenteric plexus^10^, and shared more similarity with a previous analysis of the adult myenteric plexus using single-nuclei analysis^11^. On the other hand, our analysis of the colon myenteric plexus (**Extended Data Figure 3d**) revealed less functional classes of neurons compared to previous report^11^, which analyzed a mixture of myenteric and submucosa enteric neurons. Future analysis of colon submucosa neurons is needed for a better comparison. Within the submucosal plexus of the small intestine, we identified 7 neuronal subtypes, including a population co-expressing *Chat*, *Calcb*, *Advillin* (*Avil*), and *Cysltr2*—markers of primary sensory neurons (**Extended Data Figure 4a-b**).

We presented the first comprehensive scRNA-seq atlas of the gastric enteric nervous system (ENS). Profiling 21810 gastric ENS cells revealed 2,452 enteric neurons (**Extended Data Figure 1e-f**), resolved into 10 subtypes with distinct marker genes (**Figures 1f-g**). Mirroring small intestinal and colonic myenteric neurons, *Chat* and *Nos1* expression segregated gastric myenteric neuronal subtypes. In contrast, *Slc17a6* (encoding vesicular glutamate transporter VGLUT2) was undetectable in gastric myenteric neurons but marked specific subtypes in the small intestine and colon (**Extended Data Figure 4c-e**). Notably, one gastric myenteric subtype co-expressed the immunoregulatory neuropeptide *NMU* (**Figure 1g**) and several type II cytokine receptors (**Figure 1h**), suggesting neuron-immune crosstalk in the stomach. We further identified a gastric subtype expressing *Advillin(Avil)*, *Ptgfr*, and *Piezo2*—genes characteristic of dorsal root ganglion Aβ low-threshold mechanoreceptors (Aβ-LTMRs)^35^—indicating the presence of gastric enteric mechanosensors (**Figures 1g-h, and Extended Data Figure 4c**).

The ENS works synergistically with the autonomic nervous system to regulate gastrointestinal function. Systematic analysis of cholinergic (parasympathetic) and adrenergic (sympathetic) receptor expression across ENS populations revealed plexus-specific patterns. Myenteric neurons broadly expressed muscarinic (*Chrm1*, *Chrm2*, *Chrm3*) and nicotinic (*Chrna3*, *Chrna7*) receptors, with *Chrm2* and *Chrna3* as the most abundant receptors (**Figure 1h, Extended Data Figure 5a–b**). Submucosal neurons exhibited preferential expression of nicotinic receptors (*Chrna3*) over muscarinic types (**Extended Data Figure 5c**). Adrenergic receptors included two α2-adrenergic subtypes and one β-adrenergic receptor (**Figure 1h, Extended Data Figure 5a–c**). Critically, neither cholinergic nor adrenergic receptors were significantly detected in enteric glia, defining a neuron-specific framework for autonomic modulation of gastrointestinal physiology (**Extended Data Figure 2b**).

Gut hormones and peptides, secreted by specialized chemosensory epithelial cells, play crucial roles in regulating gastrointestinal physiology, metabolism, and immunity. We identified distinct expression patterns of gut peptide receptors across enteric neuron populations (**Figure 1h, Extended Data Figure 5a-c**). Notably, myenteric and submucosal neurons express different receptor repertoires: myenteric neurons predominantly express receptors for GLP-1, GLP-2, and CCK, while submucosal neurons show preferential expression of neurotensin receptor 1 (*Ntsr1*). Receptors for PYY (*Npy1r*, *Npy2r*, *Npy4r*) are found in both myenteric plexus and submucosa plexus. These findings suggest compartmentalized regulation of enteric neural circuits by gut-derived signals. We further observed region-specific receptor distribution along the gastrointestinal tract. For example, while CCKAR is expressed in both stomach and small intestine myenteric plexuses, CCKBR appears restricted to gastric myenteric neurons (**Figure 1h**). This anatomical distribution aligns with known pharmacological properties - both receptors respond to CCK, but only CCKBR is activated by physiological concentrations of gastrin, a stomach-derived peptide that stimulates gastric acid secretion^36,37^. These expression patterns likely reflect specialized functional adaptations to local gut microenvironments.

Analysis of mechanosensitive ion channel expression revealed plexus- and region-specific patterning within the enteric nervous system (ENS). *Piezo2* dominated in gastric myenteric neurons, while *Piezo1* was the principal channel in the submucosal plexus (**Figure 1h, Extended Data Figure 5a-c**). In the small intestine and colon, *Piezo1* and *Piezo2* segregated into distinct myenteric neuron subtypes—consistent with recent functional evidence implicating *Piezo1* in enteric mechanotransduction^38^. This compartmentalized expression indicates specialized mechanosensory roles for enteric neurons across gastrointestinal regions.

### A genetic toolkit for enteric neuron subpopulations

To enable systematic characterization of enteric neuron subpopulations, we curated a suite of genetic tools based on marker genes identified through scRNA-seq (**Extended Data Figure 4**). This comprehensive toolkit includes Cre and FlpO lines targeting four functionally distinct categories of enteric neurons: (1) neurotransmitter-defined populations expressing major ENS signaling molecules; (2) neuropeptide-expressing neurons marked by *Sst-Cre*, *Vip-Cre*, *Cck-Cre*, and *Calcb-FlpO*; (3) receptor-expressing neurons responsive to gut-derived signals (*Glp1r-Cre*, *Cckar-Cre*, *Npy2r-Cre*, and *Cysltr2-Cre*); and (4) putative mechanosensitive neurons labeled by *Piezo2-Cre*.

To characterize the spatial organization and projection patterns of genetically identified enteric neurons, we visualized reporter-labeled populations using either AAV-mediated local delivery or germline-encoded fluorescent proteins (**Figures 2a-d, Extended Data Figure 6**). Our comprehensive mapping revealed that each Cre line selectively labeled distinct neuronal subpopulations with remarkable specificity along the gastrointestinal tract, enabling systematic morphological and functional characterization. Consistent with our scRNA-seq data, we observed differential targeting of enteric neuron populations: while *Sst-Cre*, *Cysltr2-Cre*, *Calcb-FlpO* and *Npy2r-Cre* labeled both myenteric and submucosal neurons (**Figures 2a-d and Extended Data Figure 6a**), other reporters (e.g., *Vglut2-Cre*, *Gad2-Cre* and *Piezo2-Cre*) were restricted to myenteric populations (**Figures 2a-d**). Detailed analysis of projection patterns revealed further specialization - some myenteric subtypes (e.g. *Piezo2-Cre*) exclusively innervate the muscularis externa, whereas others (e.g., *Gad2-Cre*) send elaborate terminals into the submucosal plexus, suggesting potential roles in coordinating responses across gut wall layers.

**Figure 2.**
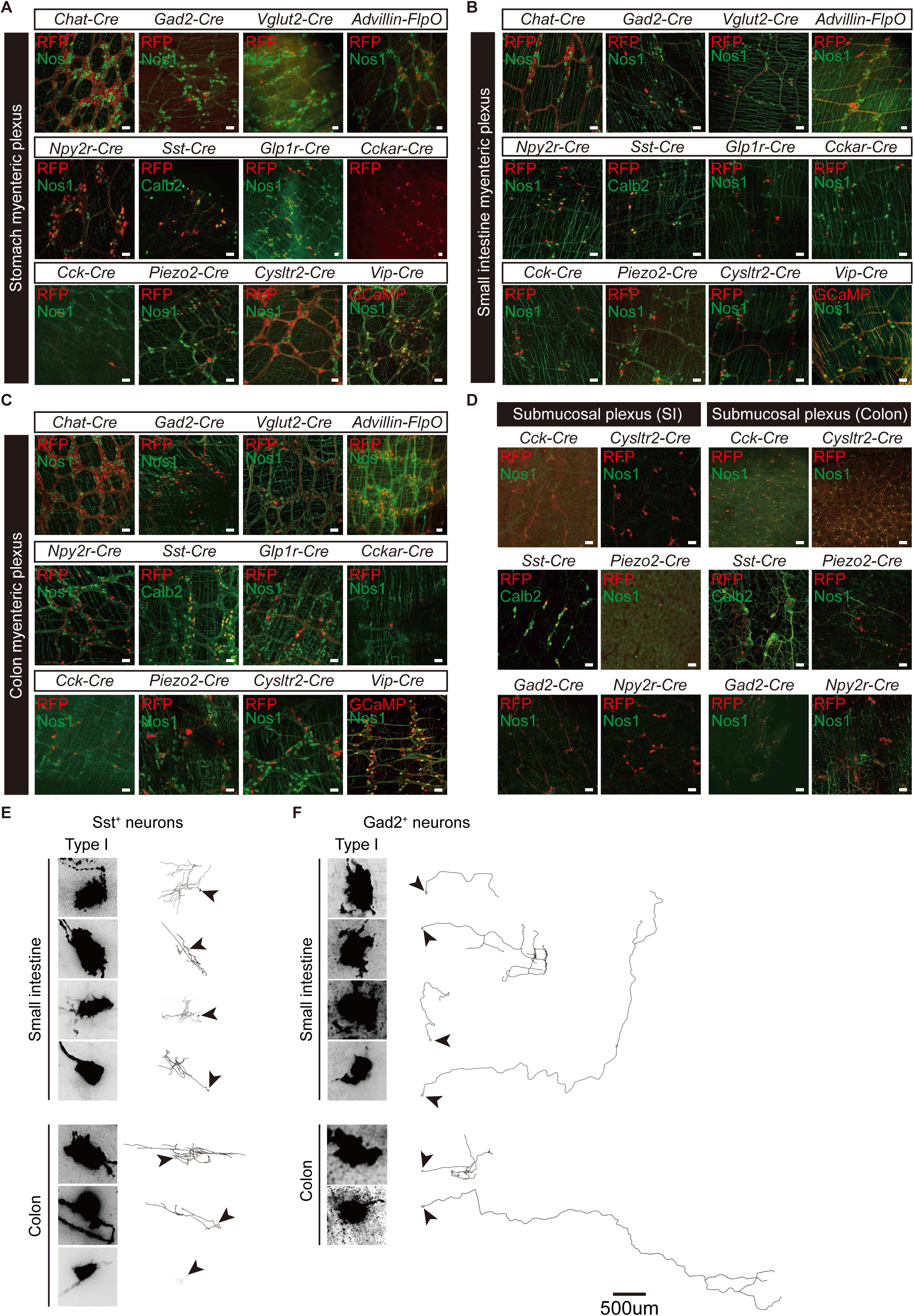
Genetic toolkits for molecularly defined enteric neuron subclasses. **(a)** Representative images for stomach myenteric neurons labeled by indicated recombinase reporter lines. Enteric neurons are labeled by immunostaining for RFP or GCaMP driven by recombinase reporter lines. Nos1 or Calb2 counterstaining is used to visualize the enteric ganglia. Scale bar, 100 μm. **(b)** Representative images for small intestine myenteric neurons labeled by indicated recombinase lines. Scale bar, 100 μm. **(c)** Representative images for colon myenteric neurons labeled by indicated recombinase lines. Scale bar, 100μm. **(d)** Representative images for submucosa enteric neurons in the small intestine (SI) and colon labeled by indicated recombinase lines. Scale bar, 100 μm. **(e-f)** Representative images of reconstructed single neuron showing the total morphology of *Sst^+^* or *Gad2^+^* myenteric neurons from the small intestine and colon. Scale bar, 500 μm.

To characterize the single cell morphology of genetically identified myenteric neurons, we performed sparse labeling through local injections of low-titer AAV2/9-hSyn-FLEX-tdTomato into the gut wall. We performed this analysis for myenteric neurons marked by *Gad2-Cre*, *Sst-Cre*, *Cck-Cre*, *Cysltr2-Cre*, *Glp1r-Cre*, *Advillin-Cre^ER^*, *Advillin-FlpO* and *Vglut2-FlpO* (**Figures 2e-f, Extended Data Figure 9 and Extended Data Figure 10**). This approach revealed distinct morphological subtypes: *Sst-Cre*-labeled neurons displayed uni-axonal morphology with extensive local branching (**Figure 2e**), while *Gad2-Cre*^+^ neurons exhibited classic Dogiel type I characteristics - thorny somatic processes and a single aboral-projecting axon with variable branching patterns (**Figure 2f**). The morphological features of *Gad2-Cre^+^* neurons, particularly their submucosal projections, reveal their identity as descending interneurons that also regulate secretomotor reflexes. Additional subtype-specific morphological analyses are described in subsequent sections (**Extended Data Figure 9 and 10**).

### Small intestine myenteric neurons are activated by nutrients, irritants and cytokine

The enteric nervous system monitors luminal contents regulate gut function^3,13,14^. To investigate what luminal signals can influence the activity of myenteric neurons, we developed an *ex vivo* calcium imaging approach^34^ (**Figure 3a**, see details in the method). We expressed the genetically encoded calcium indicator GCaMP6^39^ pan-neuronally in the enteric nervous system using either a transgenic mouse line (Snap25-GCaMP6s^40^) or an adeno-associated virus (AAV) with a neuronal promoter (AAV-hSyn-GCaMP6s). This preparation allowed administration of sensory stimuli on the mucosa side while recording spatially resolved calcium dynamics in individual myenteric neurons. In a single preparation, we can approximately image from 40 to 80 enteric neurons and quantify their responses to up to 6 consecutive stimuli. We focused our analysis of the myenteric plexus of the small intestine.

**Figure 3.**
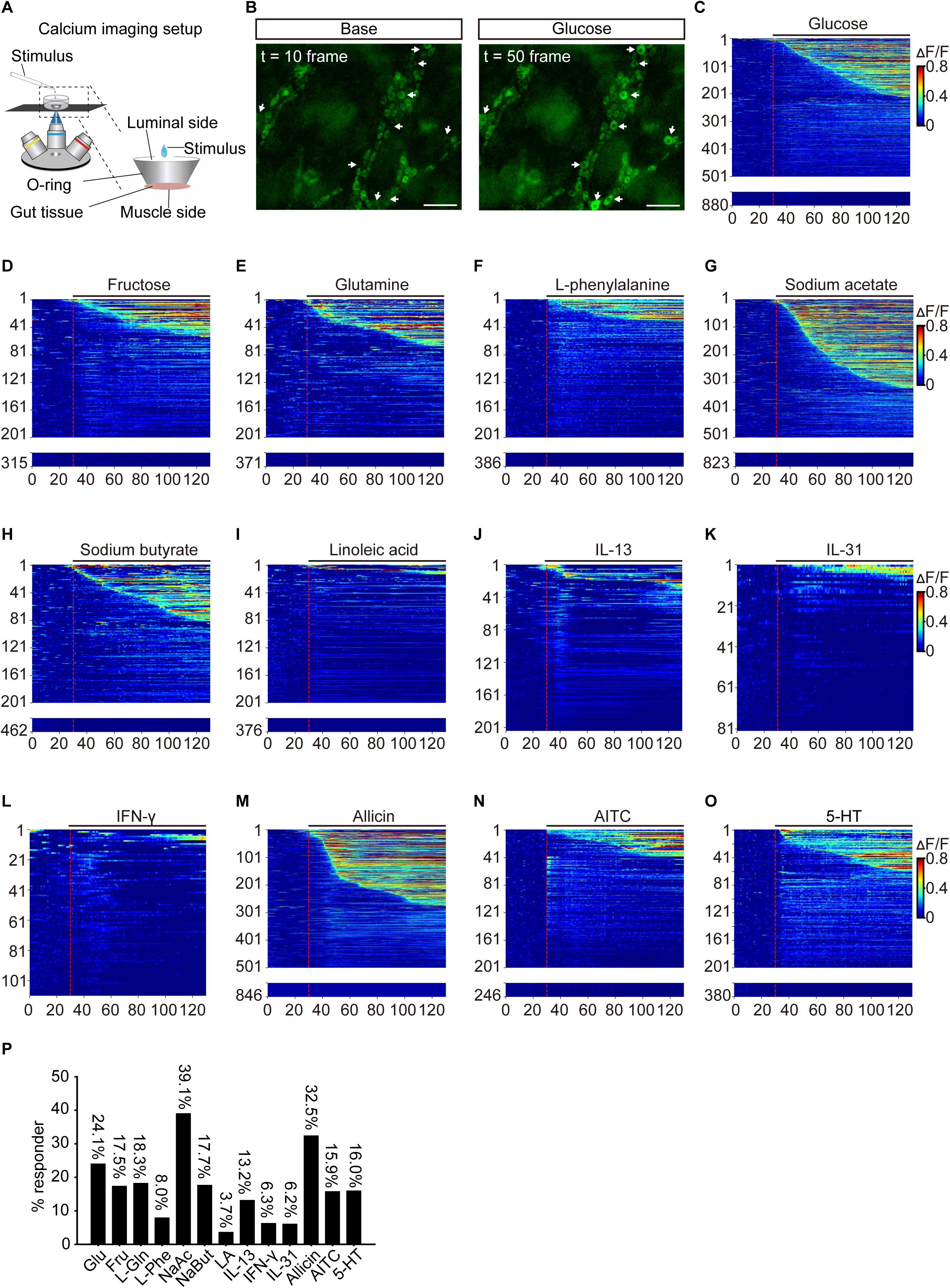
Small intestine myenteric neurons sense multiple luminal stimuli. **(a)** Schematic representation of the *ex vivo* calcium imaging setup. A custom stainless-steel O-ring was utilized to fix the small intestine tissue, ensuring that the luminal side faced upwards for acute application of chemical stimuli, while the muscular side was positioned downwards for fluorescence live imaging. **(b)** Representative images showing enteric neurons (arrowheads) respond to glucose. Before stimulation, tL=L10Lframe (left) and after stimulation, tL=L50Lframe (right). Stimuli were delivered at t = 30 frame. Scale bar, 100 μm. **(c-o)** Heatmaps depicting calcium responses (ΔF/F, color-coded) of neurons to each stimuli. A total of 880 neurons were recorded in response to 125 mM glucose (**c**), 315 neurons were recorded for 125 mM fructose (**d**), 371 neurons for 100 mM L-glutamine (**e**), 386 neurons for 100 mM L-phenylalanine (**f**), 823 neurons for 50 mM sodium acetate (**g**), 462 neurons for 50 mM sodium butyrate (**h**), 376 neurons for 100 mM linoleic acid (**i**), 204 neurons to the inflammatory cytokine 100 ng/ml IL-13 (**j**), 81 neurons to the inflammatory cytokine 100 ng/ml IL-31 (**k**), 110 neurons to 100 ng/ml IFN-γ (**l**), 846 neurons for 0.5% allicin (an activator of TRPA1 channel, **m**), 246 neurons for 1% AITC (**n**), and 380 neurons for 100 μM 5-HT (**o**). The red dashed line in the heatmap indicates the time point of stimuli application. **(p)** Quantification for proportion of responders to indicated stimuli.

First, we examined the effects of nutrients on enteric neuronal activity. We applied chemicals to the mucosal side of the preparation as nutrients are restricted within the gut lumen under physiological conditions (**Figure 3a**). Glucose, a key nutrient detected by gut-innervating sensory neurons, was tested at concentrations ranging from 50 mM to 300 mM. We observed a dose-dependent activation of myenteric neurons, with higher glucose concentrations eliciting stronger responses (**Extended Data Figure 7a**).

Next, we expanded our survey to multiple nutrient-related chemicals, including two simple sugars (glucose and fructose), two short-chain fatty acids (sodium acetate and sodium butyrate), one long-chain fatty acid (linoleic acid), and two amino acids (L-glutamine and L-phenylalanine). We observed varying degrees of neuronal activation across these stimuli: glucose and fructose activated 24.1% and 17.5% of myenteric neurons (**Figures 3b-d, and Figure 3p**) respectively, while L-glutamine and L-phenylalanine activated 18.3% and 8.0% (**Figures 3e-f, and Figure 3p**) respectively. Fatty acids also activated myenteric neurons, with short-chain fatty acids (17.7% for sodium butyrate, 39.1% for sodium acetate) inducing stronger responses than the long-chain fatty acid linoleic acid (3.7%) (**Figures 3g-i, and Figure 3p**).

Given the close anatomical and functional associations between enteric neurons and immune cells^41,42^, we examined the expression of cytokine receptors in our single-cell RNA-seq data. We found that multiple receptors for cytokines (e.g. *Il4ra*, *Il13ra1*, *Il31ra*, and *Ifngr1/Ifngr2*, **Extended Data Figure 5a**) were expressed in specific subsets of enteric neurons, including putative intrinsic primary afferent neurons (IPANs; see below for definition)^43,44^. Notably, one IPAN subset expressed the neuropeptide NMU, a known regulator of type 2 innate lymphoid cells^45,46^. This prompted us to investigate whether myenteric neurons respond to these cytokines. We stimulate the tissue with IL-13, IL-31, or IFN-γ from the mucosal side, and observed that 13.2%, 6.2%, and 6.3% of myenteric neurons were activated by these cytokines respectively (**Figures 3j-l and Figure 3p**).

We also tested whether myenteric neurons respond to chemical irritants such as allyl isothiocyanate (AITC) and allicin, plant-derived compounds that activate the TRPA1 cation channel and cause gastrointestinal irritation upon ingestion^47,48^. When administrated from the luminal side, approximately 15.9% of myenteric neurons responded to luminal AITC, while 32.5% responded to allicin (**Figures 3m-n and Figure 3p**). Notably, we did not detect *TRPA1* transcripts in enteric neurons (**Extended Data Figure 5a**), suggesting that the observed responses are not due to direct activation of myenteric neurons. Previous studies reported that both AITC and allicin can activate enterochromaffin cells. Therefore, our results indicate an enterochromaffin-neuron crosstalk (see below).

Next, we tested whether different nutrients activate distinct subsets of enteric neurons in the myenteric plexus. To address this, we applied 5-6 different stimuli consecutively to the same imaging preparation and analyzed the evoked neuronal responses (**Figures 4a-b**). We examined approximately 1700 neurons that responded to at least one of the 6 stimuli tested and categorized their tuning properties. A large proportion of myenteric neurons are activated by more than one nutrient (**Figure 4c**). To quantify the response properties, we grouped neurons based on their sensitivity to a specific nutrient (e.g. neurons responding to glucose were grouped together and termed “glucose-responsive neurons”) and quantified the proportion of singly-tuned responders and multi-tuned responders among different groups. Around 10%-40% of neurons responded exclusively to one stimulus (singly-tuned responder), while 60%-90% responded to multiple stimuli (multi-tuned responder). To systematically evaluate the overlap in neuronal responses, we calculated a convergence index (see Methods) for each pair of stimuli. Our analysis suggested variable degrees of convergence between different nutrients. In general, fructose and linoleic acid showed less degree of convergence with other nutrients. Interestingly, glucose and short-chain fatty acids have higher degrees of convergence compared to glucose and fructose (**Figure 4d**).

**Figure 4.**
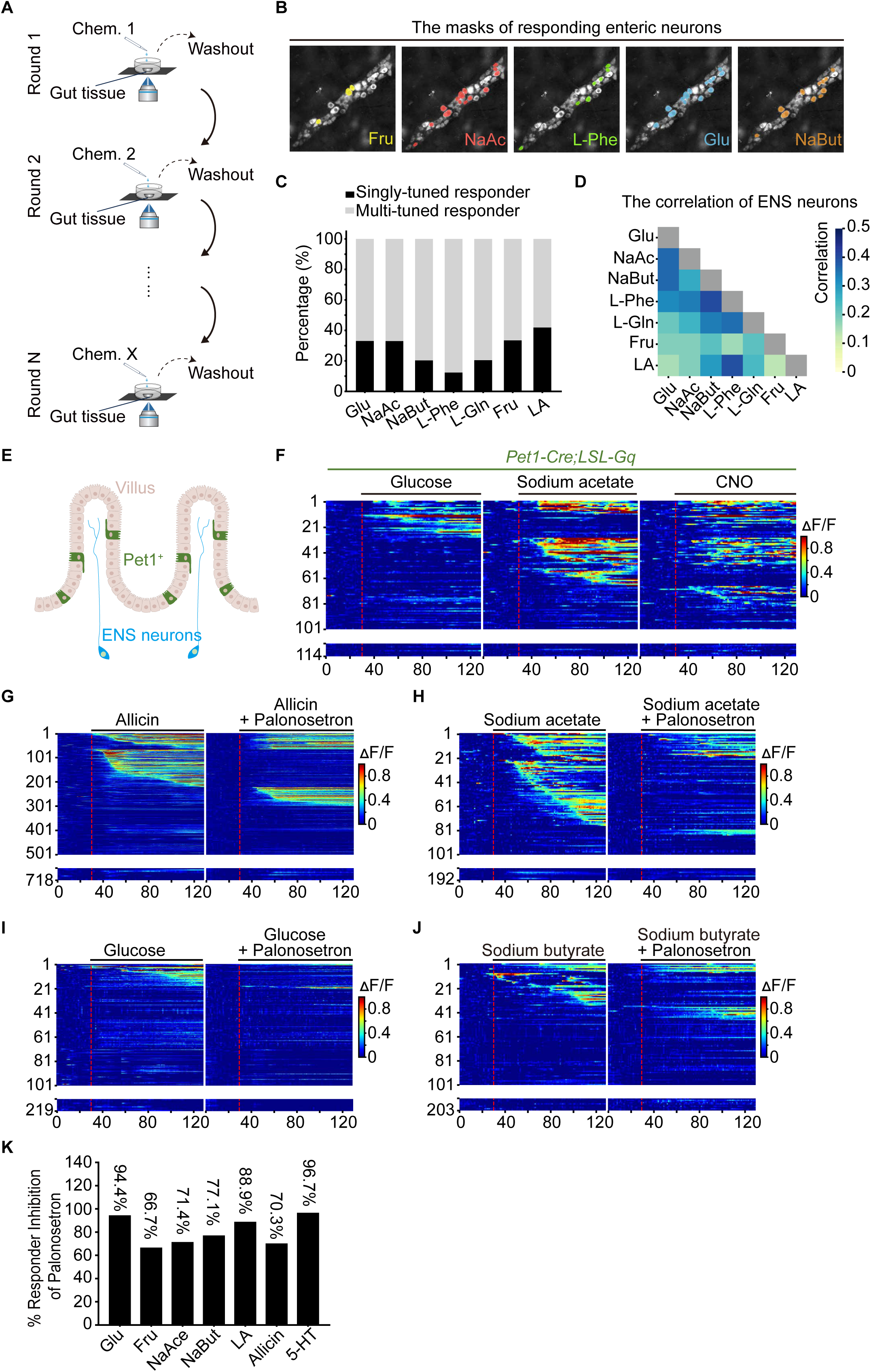
Mucosa-enteric pathway mediates myenteric responses to luminal nutrients. **(a)** Schematic diagram of multi-round *ex vivo* calcium imaging **(b)** The representation images and masks indicate the responding myenteric neurons to indicated stimuli. **(c)** The graph illustrates the proportions of singly-tuned responder (black bars) and multi-tuned responder neurons (gray bars) among myenteric neurons activated by indicated stimuli. **(d)** Correlation analysis of neurons’ responses to two different stimuli annotated in the bottom and left of the checkboard. The color intensity represents the correlation coefficient. **(e)** A cartoon illustration of the crosstalk between enteric neurons and *Pet1^+^* enterochromaffin cells. **(f)** Heatmaps depicting calcium responses (ΔF/F) of myenteric neurons following chemogenetic activation enterochromaffin cells. Notably, many of these activated neurons is also responsive to glucose or sodium acetate. The red dashed line in the heatmap indicates the administration of the stimuli. Each row in the heat maps indicates the same neuron. **(g-j)** Heatmaps depicting calcium responses (ΔF/F) of myenteric neurons to 0.5% allicin (**g**), 50 mM sodium acetate (**h**), 125 mM glucose (**i**), and 50 mM sodium butyrate (**j**) before and after 1 μM Palonosetron (an antagonist of 5-HT_3_R). The red dashed line in the heatmap indicates the administration of the stimuli. Each row in the two heat maps from the same panel indicates the same neuron. Palonosetron were incubated for the luminal side 5 min before recording and were maintained throughout the recording session. **(k)** Quantification for proportion of responding neurons inhibited by Palonosetron.

### Epithelial—enteric neuron signaling contributes to nutrient sensing

The small intestinal epithelium harbors specialized chemosensory cells, such as enteroendocrine/enterochromaffin cells^15^. These chemosensory cells express a diverse array of surface receptors, including G protein-coupled receptors (GPCRs) and ion channels, enabling them to detect and respond to a wide range of luminal stimuli such as sugars, fatty acids, and amino acids^49,50^. Upon chemical stimulation, they release signaling molecules such as ATP, 5-HT and various gut peptides, which can activate nerve terminals reaching the villi (**Figure 4e**). Using our imaging preparation, we recorded 5-HT evoked myenteric neuron response (**Figure 3o**). To directly test functional coupling between chemosensory epithelium and enteric neurons, we expressed chemogenetic activator hM3Dq^51^ in 5-HT-positive enteroendocrine/enterochromaffin cells using *Pet1-Cre* (**Extended Data Figure 7b** for *pet1-Cre* expression pattern), activated them with CNO and monitored the enteric neuronal activity with GCaMP6s. Consistent with the existence of a functional coupling, we observed robust activation of enteric neurons upon CNO administration (**Figure 4f**). Interestingly, a large proportion of the CNO-responding neurons are also activated by luminal nutrients such as glucose or sodium acetate.

*Htr3a and P2rx2* are the predominant serotonin and ATP receptors expressed in myenteric neurons (**Extended Data Figure 5a**), and HTR3 inhibitor palonosetron can inhibit more than 90% of 5-HT induced myenteric response (**Figure 4k, and Extended Data Figure 7c**). Therefore, we tested their contribution to nutrient induced responses. Blocking HTR3 using Palonosetron diminished a large proportion of nutrient induced responses in myenteric neurons (**Figures 4g-j, and Extended Data Figure 7d-e**). On the other hand, blocking P2X receptor by Gefapixant led to a smaller degree of reduction in nutrient induced responses (**Extended Data Figure 7f-j**). Comparing the inhibitory effect of these two inhibitors, it appears that glucose responses are predominantly mediated by 5-HT.

### Genetic access to putative enteric sensory neurons

Next, we develop tools for genetic access to enteric sensory neurons. Among the genetic markers screened, we identified enteric neurons expressing *Advillin*, a highly specific marker widely used to label dorsal root ganglion (DRG) and vagal sensory neurons^21^. Single-cell RNA sequencing revealed 3-4 subsets of *Advillin*^+^ myenteric neurons, as well as a subset of *Advillin*-expressing submucosal neurons in the small intestine (**Figures 5A-D, Figures S8A-B**). Many of these neurons co-expressed *Calcb*, encoding CGRPβ—a classic neuropeptide associated with enteric sensory neurons (**Figures S4B-E and Figures S8B-E**)—along with multiple receptors for signaling molecules derived from chemosensitive enterochromaffin and enteroendocrine cells, indicating these neurons’ ability to sense luminal chemicals. Furthermore, a subset of *Advillin*-positive myenteric neurons expresses mechanosensitive channel Piezo2 (**Figures S8B-E**). Based on these findings, we propose that *Advillin* marks intrinsic primary afferent neurons (IPANs) in the murine enteric nervous system.

**Figure 5.**
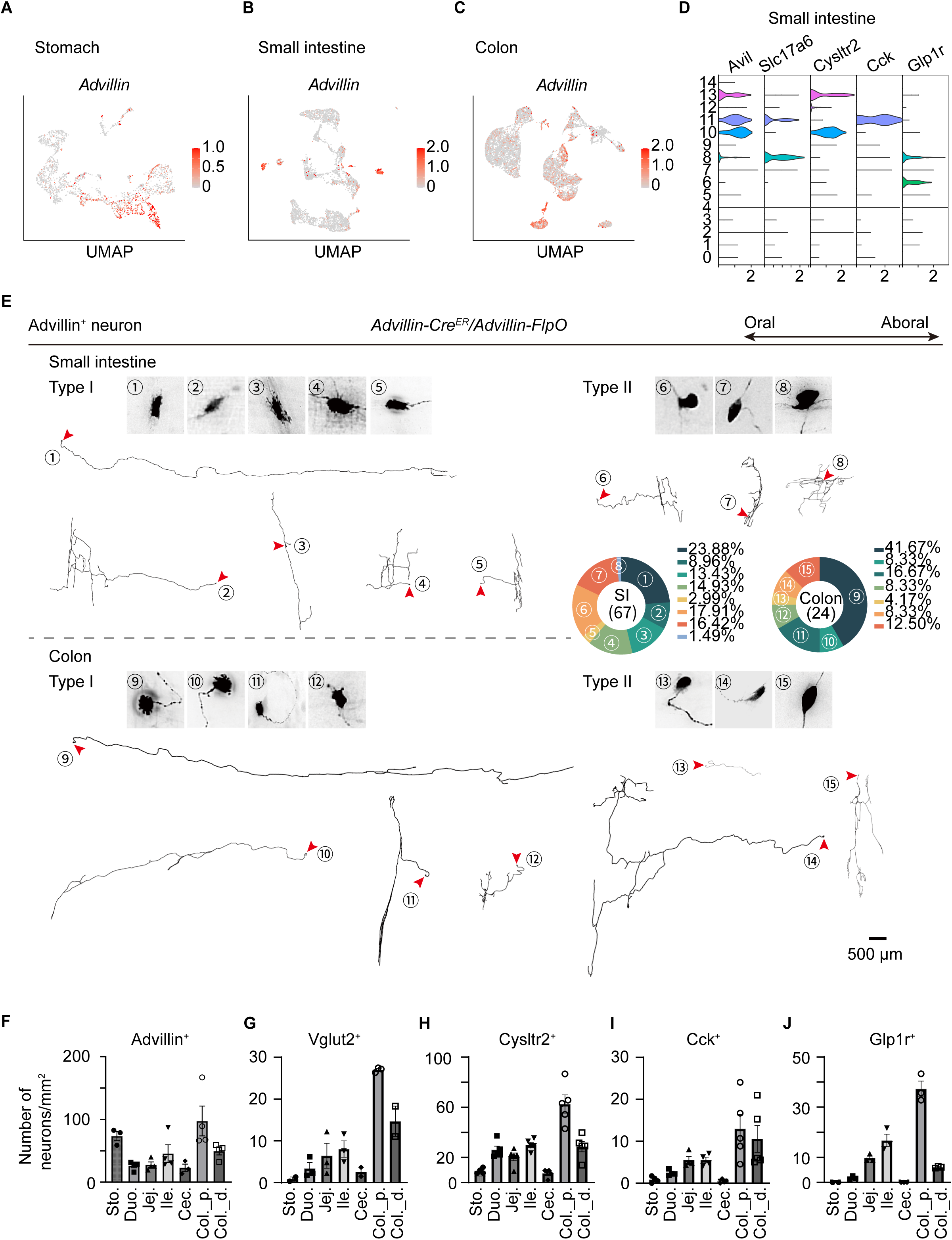
Genetic access to putative intrinsic primary afferent neurons. (a-c) Feature plot showing the expression of *Advillin* in myenteric plexus of the stomach (**a**), small intestine (**b**), colon (**c**). **(d)** Volin plot showing the expression of indicated marker gene in small intestine. **(e)** Reconstructed single neuron morphology of *Advillin^+^* myenteric neurons from the small intestine and colon. A total of 91 neurons with 15 morphological types (divided into Dogiel type I and Dogiel type II) were analyzed by sparse labeling. The distribution of each morphological type shown is quantified in small intestine and colon separately and shown in the pie chart. Scale bar, 500 µm. **(f-j)** Bar graph showed that quantification of indicated IPANs subtype within the gastrointestinal tract. Error bars represent mean ± SEM. Each dot indicated quantification from one animal. Data from 3 animals were used for quantification.

To genetically target *Advillin*^+^ enteric neurons, we used *Advillin-Cre^ER^* or *Advillin-FlpO* mice in combination with Cre- or FlpO-dependent reporters. Both reporters robustly labeled myenteric neurons throughout the gastrointestinal tract, as well as neurons in the submucosal plexus of the small intestine and colon (**Figures 2A-C, Figures S6B-C and Figure S9**). Notably, nearly all *Advillin*^+^ neurons were Nos1-negative and lacked projections into the smooth muscle layers of the muscularis externa, supporting their classification as intrinsic primary afferent neurons (**Figure S6C**).

To characterize the morphological diversity of *Advillin*^+^ myenteric neurons, we performed sparse labeling and analyzed 67 small intestinal and 24 colonic neurons (**Figure 5E**). These neurons exhibited 15 distinct morphological subtypes, with approximately 25-35% displaying classical Dogiel type II characteristics LJ multiple axons emerging from a smooth soma and relatively confined innervation fields within the myenteric plexus. In contrast, the majority of Dogiel type I neurons featured axonal projections extending far from their soma, though a subset maintained more localized axonal arbors.

*Advillin^+^* enteric neurons comprise 3-4 molecularly distinct subclasses. Mining our small intestine scRNA-seq data revealed *Cck*, *Glp1r*, and *Cysltr2* as markers defining different subsets of *Advillin^+^* myenteric neurons in the small intestine, two of which also expressed *Slc17a6* (encoding Vglut2, **Figure 5D**). To selectively characterize these populations, we employed intersectional genetic strategies LJ particularly crucial since both *Cck-Cre* and *Glp1r-Cre* exhibit broader expression in other GI cell types (data not shown). By combining *Vglut2-FlpO* with either *Glp1r-Cre* or *Cck-Cre* and the dual-recombinase reporter *Ai65*^40^, we achieved specific tdTomato expression in *Glp1r^+^* or *Cck^+^* myenteric neurons (**Figure 5D, Figure S9**), respectively. For *Cysltr2^+^*, we used *Phox2b-FlpO* and *Cysltr2-Cre* with an alternative dual-recombinase reporter to drive GFP expression exclusively in FlpO^+^/Cre^+^ cells (**Figure 5D, Figure S9**).

These intersectional genetic approaches enabled comprehensive mapping of neuronal distributions along the gastrointestinal tract (**Figures 5F-G, and Figure S9**). *Advillin*^+^ myenteric neurons showed highest density in the stomach and proximal colon compared to the regions. However, *Vglut2, Glp1r or Vglut2, Cck-*double positive neurons were scarce in the stomach but progressively increased in density toward the distal small intestine. On the other hand, *Cysltr2*^+^ neurons maintained a relatively uniform distribution throughout the small intestine. These anatomical features may suggest regional specifications of IPANs.

Next, we performed single neuron morphological analysis using *Vglut2* (**Figure S10A**), *Cysltr2* (**Figure S10B**), *Cck* (**Figure S11A**), and *Glp1r* reporters (**Figure S11B**). More than a third of *Cysltr2*^+^ myenteric neurons belonged to Dogiel type II. Notably, a large proportion of *Cysltr2-Cre*-labeled neurons displayed intricate local projection patterns, while *Vglut2^+^* neurons typically projected aborally with variably branched terminal arbors. *Glp1r-Cre*, which labeled two small intestine myenteric subpopulations (one is *Vglut2^+^*and the other is *Gad2*^+^), labeled neurons with aboral long-range projecting axons. On the other hand, *Cck-Cre*, which labeled a different Vglut2-positive myenteric neurons in the small intestine, displayed a greater variety of morphology. Collectively, our intersectional genetic strategies combined with viral labeling and sparse tracing revealed comprehensive anatomical and morphological features of putative sensory neurons within the myenteric plexus.

### Enteric sensory neurons participate in sensing luminal signals

Next, we characterized the response profiles of enteric sensory neurons to diverse luminal stimuli. We generated *Advillin-Cre^ER^;*Ai162 mice to express GCaMP6s in *Advillin^+^* enteric neurons^52^ (**Figure 6a)** and performed *ex vivo* calcium imaging in the small intestine using our established preparation (**Figure 3a)**. This enabled us to capture the response patterns of *Advillin^+^*neurons to sequential luminal delivery of 6–7 stimuli within the same field of view (**Figure 6b**).

**Figure 6.**
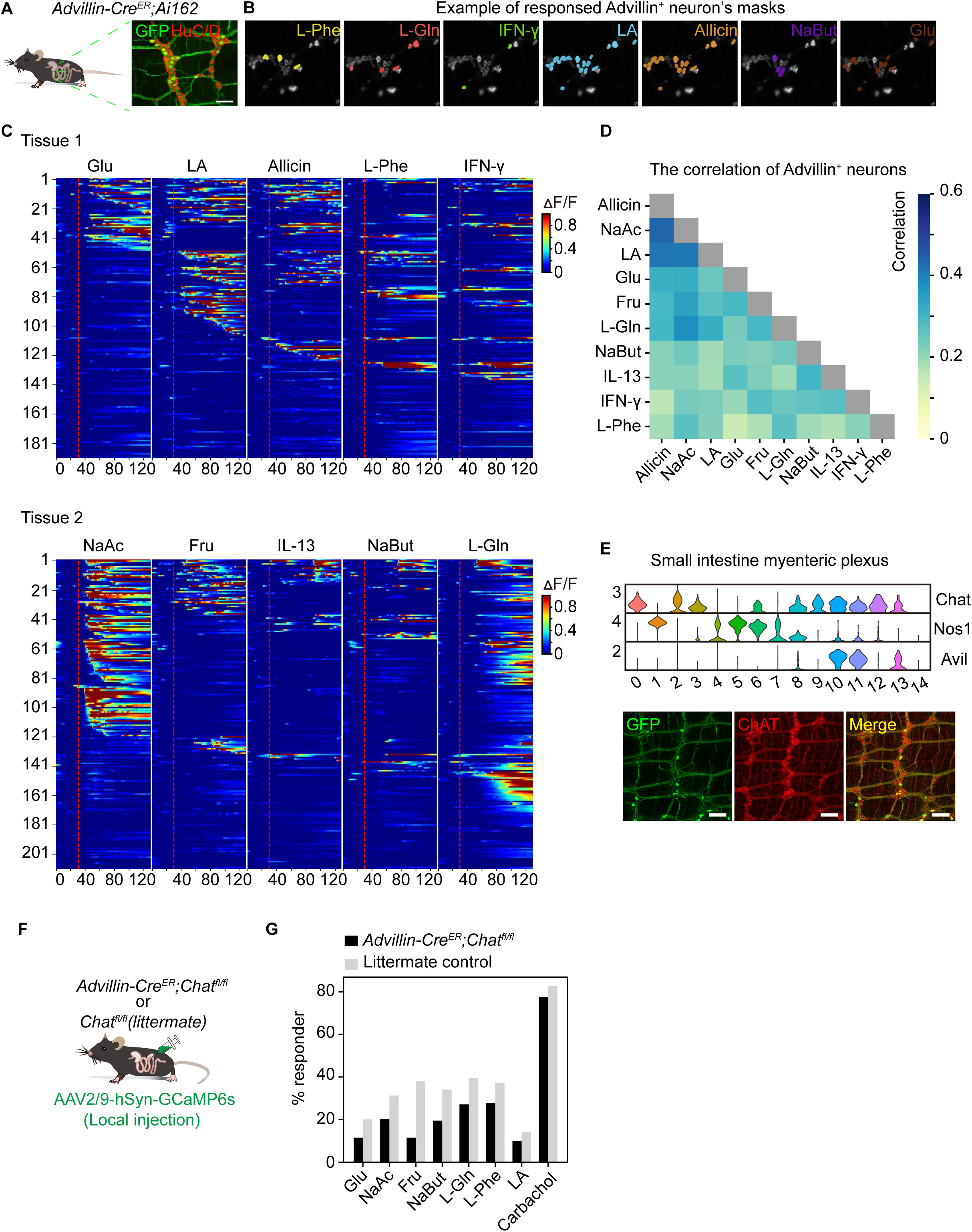
Sensory responses of *Advillin^+^* enteric neurons. **(a)** Schematic and immunofluorescent stanning of *Advillin^+^* enteric neuron expression GCaMP6. HuC/D is used as counterstaining for all myenteric neurons. Scale bar, 100 μm. **(b)** The representation images and masks of the responding neurons to indicated stimuli. **(c)** Heatmaps depicting calcium responses (ΔF/F, colour-coded) of *Advillin^+^* enteric neurons to different stimuli. Each row from five heat maps in the same tissue represents the response from the same neuron to five different stimuli. Stimuli for the tissue 1 in the upper panel are: glucose, linoleic acid, allicin, L-phenylalanine, IFN-γ. Stimuli for the tissue 2 in the lower panel are: sodium acetate, fructose, IL-13, sodium butyrate, and L-glutamine. The red dashed line in the heatmap indicates the time point of stimuli administration. **(d)** Correlation analysis of *Advillin^+^* neuron’s responses to two different stimuli annotated in the bottom and left of the checkboard. The color intensity represents the correlation coefficient. **(e)** Violin plot and immunostaining showing the co-expression of *Advillin* and *Chat*. GCaMP immunostaining is performed in *Advillin-Cre^ER^*;Ai162 mice to label *Advillin^+^* neurons. **(f)** A cartoon schematic illustrates the local administration of AAV2/9-hSyn-GCaMP6s in the small intestine of *Advillin-Cre^ER+/-^;Chat^fl/fl^ or Chat^fl/fl^* mice for *ex vivo* calcium imaging. **(g)** Quantification of the proportion of myenteric neurons in the small intestine that respond to indicated stimuli in *Advillin-Cre^ER+/-^;Chat^fl/fl^*or *Chat^fl/fl^* mice (littermate control) animals. Carbachol was used as a positive control to activate enteric neurons.

Across more than 700 *Advillin^+^* neurons, we identified distinct chemical-specific activation profiles: most neurons exhibited multi-tuned responses to diverse stimuli, confirming polymodal coding logic in the myenteric plexus (**Figure 6c and Figure 4c**). Correlation analysis revealed hierarchical nutrient encoding LJ for instance, responses to glucose (Glu) and sodium acetate (NaAc) were strongly correlated (r² = 0.4), whereas glucose (Glu) and IFNγ showed minimal correlation (r² < 0.1; **Figure 6d**). Notably, allicin responses correlated with multiple nutrients (r² > 0.3), suggesting enterochromaffin (EC) cells mediate luminal chemosensation—consistent with our chemogenetic activation data (**Figure 4e-f**).

Enteric sensory neurons transmit sensory induced neuronal response to the result of the enteric neuronal network. To block this process, we attenuated neurotransmitter release from enteric sensory neurons. Given the pivotal role of acetylcholine (ACh) in enteric signaling, we confirmed ubiquitous ChAT co-expression in *Advillin^+^* neurons via scRNA-seq and immunofluorescence (**Figure 6e**). To functionally interrogate this pathway, we generated *Advillin-Cre^ER^*;*Chat^fl/fl^* mice, selectively ablating ACh synthesis in *Advillin^+^* neurons. Abolishing ACh synthesis led to a reduction in nutrient-evoked responses among myenteric neurons (**Figure 6e-g**), establishing *Advillin^+^* IPANs as essential transducers or amplifiers of luminal nutrient signals to other myenteric neurons. Nevertheless, a substantial number of myenteric neurons retained their response to luminal nutrients, consistent with model that IPANs sense luminal nutrients through chemosensory epithelium (independent of ACh release from other enteric neurons).

### Optogenetic activation of enteric neurons modulates gut motility

The ENS operates as a self-contained neural network capable of autonomous control over gastrointestinal function^2,3^. To directly interrogate how specific enteric neuron subpopulations regulate gut motility, we developed a targeted optogenetic strategy. We achieved cell-type-specific expression of the light-sensitive cation channel through two complementary approaches: (1) crossing Cre-driver mice with the Cre-dependent ChR2 reporter line Ai32^53^ (**Extended Data Figure 12g**), or (2) using dual-recombinase control by mating Cre/FlpO mice with the conditional CatCh^54^ (also a light activated cation channel) allele (Ai80) for intersectional targeting (**Figure 7a, Extended Data Figure 12h-l**). This genetic framework enabled precise optical control of defined enteric neuron populations while preserving their native circuit connectivity.

**Figure 7.**
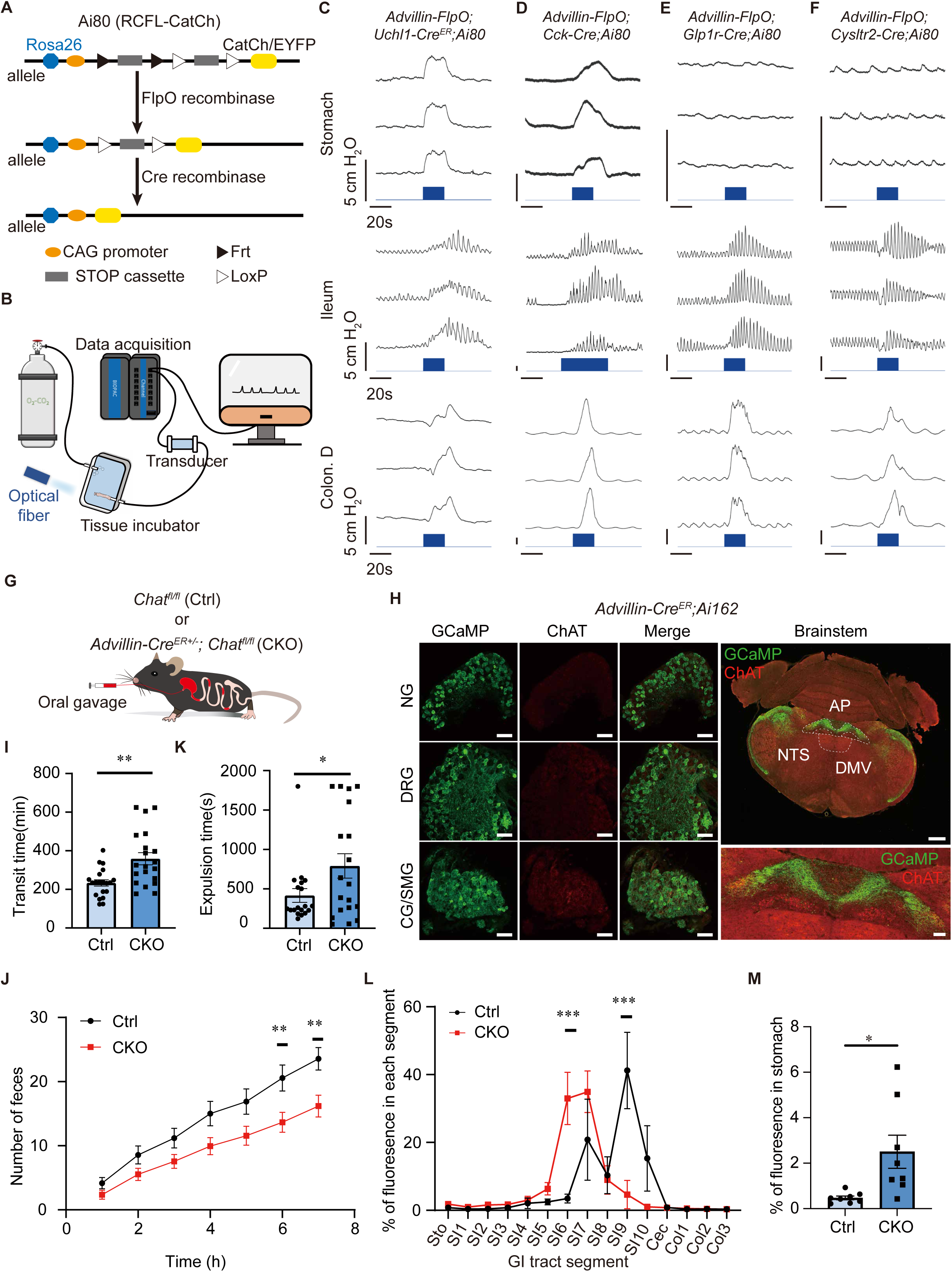
Differential regulation of gut motility by *Advillin^+^* enteric neurons. **(a)** Schematic illustration for Opsin expression in *Advillin^+^* subpopulations by intersectional genetics. **(b)** Experimental setup for optogenetic control of gut motility. **(c-f)** Representative traces showing that optogenetic induced gut motility change. Gut motility change is reflected by alteration in luminal pressure (centimeter H_2_O pressure) measured by the pressure sensor. Blue bar in the bottom indicates laser stimulation. **(g)** Schematic diagram for total GI transit assay. **(h)** Expression pattern of *Advillin-Cre^ER^* (staining for GCaMP6 in *Advillin-Cre^ER^;*Ai162 animal) in gut-related sensory neurons and autonomic motor neurons. *Advillin-Cre^ER^* is widely expressed in sensory neurons in the nodose ganglia and DRG (Chat-negative) but not Chat-positive DMV motor neurons. Coronal brain sections of the entire brain scale bar = 500 μm, other sections scale bar = 100 μm. **(i)** Total GI transit time of littermate control animals (*Advillin-Cre^ER-/-^;Chat^fl/fl^*, n = 20) and CKO animals (*Advillin-Cre^ER+/-^;Chat^fl/fl^*, n = 20), unpaired two-tailed t test: **p < 0.01. Data are expressed as mean ± SEM. **(j)** The number of feces expelled over 7 hours by littermate control animals (*Advillin-Cre^ER-/-^;Chat^fl/fl^*, n = 20) or CKO animals (*Advillin-Cre^ER+/-^;Chat^fl/fl^*, n = 20), Two-Way ANOVA: **p < 0.01. Data are expressed as mean ± SEM. **(k)** Distal colonic transit time measured as bead expulsion time in littermate control animals (*Advillin-Cre^ER-/-^;Chat^fl/fl^*, n = 19) and CKO animals (*Advillin-Cre^ER+/-^;Chat^fl/fl^*, n = 19), unpaired two-tailed t test: *p < 0.05. Data are expressed as mean ± SEM. **(l)** Quantification of the relative abundance of the FITC various segments of the GI tact after 60 minutes of oral gavage of FITC-Dextran dye in littermate control animals (*Advillin-Cre^ER-/-^;Chat^fl/fl^*, n = 6) and CKO animals (*Advillin-Cre^ER+/-^;Chat^fl/fl^*, n = 6), Data are presented as mean ± SEM, Two-way ANOVA: ***p < 0.001 **(m)** Quantification of the relative abundance of the FITC retained in the stomach after 30 minute of oral gavage of FITC-Dextran dye in littermate control animals (*Advillin-Cre^ER-/-^;Chat^fl/fl^*, n = 8) and CKO animals (*Advillin-Cre^ER+/-^;Chat^fl/fl^*, n = 8). Data are presented as mean ± SEM, *p < 0.05, unpaired t test.

To investigate how specific ENS populations regulate gastrointestinal motility, we established an *ex vivo* gut motility assay. Segments from three anatomically distinct regions (stomach, small intestine, or distal colon) were maintained in oxygenated organ chambers (**Figure 7b**) while we monitored optogenetically-induced luminal pressure changes. We first validated this approach by targeting two well-characterized enteric neuron populations: excitatory cholinergic neurons and inhibitory nitrergic neurons. As predicted, optogenetic activation of cholinergic neurons produced robust contractions, manifested as increased luminal pressure in all gut segments examined (**Extended Data Figure 12a**). Conversely, stimulation of nitrergic neurons induced relaxation, significantly reducing luminal pressure in both the stomach and small intestine (**Extended Data Figure 12b**). These results both validate our experimental system and demonstrate the opposing yet coordinated regulation of gut motility by distinct ENS subpopulations.

We extended this optogenetic approach to characterize other myenteric neuron subpopulations, revealing distinct segment-specific motility patterns: *Vglut2^+^* neurons (**Extended Data Figure 12c**) showed no gastric effect but induced transient relaxation in the ileum and a biphasic distal colonic response (transient relaxation followed by contraction); *Gad2^+^* neurons exhibited pan-gastrointestinal modulation, promoting gastric and distal colonic contractions while eliciting ileal relaxation (**Extended Data Figure 12d**); and *Cckar^+^* and *Npy2r^+^* neurons (**Extended Data Figure 12e-f**), likely regulated by endogenous gut peptides, influenced motility throughout all segments examined (stomach, small intestine, and colon).

To determine whether *Advillin^+^* myenteric neurons - putative intrinsic primary afferent neurons (IPANs) - regulate gut motility, we employed an intersectional genetic strategy. By crossing *Advillin-FlpO*; *RCFL-CatCh* mice with *Uchl1-Cre^ER^* drivers, we achieved selective optogenetic control of this population. Following tamoxifen induction, we assessed motility responses in our *ex vivo* gut preparation. Optogenetic stimulation of *Advillin^+^* neurons consistently modulated motility throughout the gastrointestinal tract (**Figure 7c**).

To further resolve functional heterogeneity among intrinsic primary afferent neurons (IPANs), we generated three distinct mouse lines targeting molecularly-defined subpopulations by crossing *Advillin-FlpO; RCFL-CatCh* mice with *Cysltr2-Cre*, *Glp1r-Cre*, or *Cck-Cre* drivers, respectively. Following validation of CatCh expression in each population (**Extended Data Figure 12j-l**), we performed optogenetic stimulation across gastrointestinal segments (**Figure 7d-f**). All three subpopulations (*Cysltr2^+^, Glp1r^+^,* and *Cck^+^*) modulated motility in both ileum and distal colon. Strikingly, only *Cck^+^*neurons elicited gastric contractions, revealing a unique role for this subset in stomach motility regulation. These results demonstrate that enteric sensory neurons can directly orchestrate gut physiology through local neural circuits, independent of central nervous system input.

### Enteric sensory neurons are critical for gut motility

To establish the functional necessity of enteric sensory neurons in gastrointestinal motility, we generated *Advillin-Cre^ER^*^+/-^; *Chat^flox/flox^* conditional knockout mice (**Figure 7G**), selectively ablating choline acetyltransferase (ChAT) - the key enzyme for acetylcholine synthesis – in *Advillin*^+^ neurons. While *Advillin-Cre^ER^* targets other peripheral sensory neurons (vagal, DRG), only enteric sensory neurons predominantly rely on acetylcholine neurotransmission (**Figure 7h**), ensuring specific disruption of enteric neural circuits.

To evaluate the effect of *Chat* deletion on gastrointestinal (GI) motility, we first measured total GI transit time by administering carmine red, a non-absorbable dye, via oral gavage and recording the time until its appearance in fecal pellets (**Figures 7g and 7i**). *Chat* conditional knockout (CKO) mice exhibited a significantly prolonged total GI transit time (233.5 ± 16.43 min) compared to littermate controls (358.6 ± 31.78 min), along with reduced fecal pellet output (**Figure 7j**). To assess distal colon motility, we performed a glass bead expulsion test, inserting a bead (2 cm from the anus) and measuring the time to expulsion. *Chat* CKO mice showed a significant delay in bead expulsion (**Figure 7k**), indicating impaired motility in the distal colon.

For upper GI motility (stomach and small intestine), we gavaged mice with FITC-Dextran and quantified the remaining fluorescence in the stomach and intestinal segments afterwards. *Chat* CKO mice retained more dye in the stomach after 30 minutes of dye gavage and exhibited slower small intestinal transit (**Figures 7l and 7m**). Together, these findings demonstrate that *Chat* expression in *Advillin*ϑ enteric neurons play a critical role in regulating gut motility.

## Discussion

The enteric nervous system is a critical hub for brain-gut axis. Defining the molecular and functional diversity is fundamental for understanding autonomic regulation of gastrointestinal physiology and gut interoception. Here, we described scRNA-seq pipelines, genetic toolkits, and neurophysiological approaches to dissect the cellular, molecular and functional diversity of the mouse enteric nervous system. These datasets and experimental approaches should empower future studies of the structure and function of the mammalian enteric nervous system with.

### Towards a comprehensive atlas of the mouse enteric nervous system

Despite the challenges to study the enteric nervous system at single cell resolution, recent advances in single cell RNA-sequencing technologies have enabled molecular analysis of the enteric nervous system at unprecedented detail^10–12^. Nevertheless, these studies are limited by using reporter lines based of different mark genes (some of which also label cells outside of the enteric nervous system), focused on different and limited segments of the gastrointestinal segments, and employed different dissociation methods. We established a customed pipeline to profile the enteric nervous system from specific segments of the gastrointestinal tract (**Figure 1**).

In our approach, we relied on reporter animals generated from a single marker gene *Phox2b*, which specifically labeled the entirety of the enteric nervous system (including both the enteric neurons and enteric glia). The advantage of using *Phox2b*-based reporters is that it can efficiently profile both the enteric neurons and enteric glia from the same animal, enable comparison of the transcriptomic changes of these two major cell types in the same condition. Our studies also demonstrated the necessity of choosing suitable fluorescent reporters: because cytosolic reporters also labeled fiber bundles and form fluorescent cellular debris contaminating the FACS sorting of enteric cells, especially for the colon and stomach whose inter-ganglionic fiber bundles are much thicker. In addition, we described a pan-neuronal marker (*Uchl1-Cre^ER^*) to enrich enteric neurons. This line is useful in experimental conditions where enrichment of enteric neurons is needed (for example, when a harsh dissociation condition is used, a disproportional enteric neuronal death would take place).

Our analysis encompassed most of the gastrointestinal tract, including the first single-cell atlas of the gastric myenteric plexus. Gastric physiology is tightly regulated by the autonomic nervous system through distinct neuronal reflexes. Previous studies have revealed labeled-line connections between vagal motor neurons and gastric enteric neurons^55^. Vagal innervation of the stomach is known to mediate multiple functions—including receptive relaxation, gastric secretion, and gastric emptying—likely through distinct subsets of vagal motor units. We hypothesize that these reflexes may be orchestrated by specialized populations of gastric enteric neurons acting downstream of different vagal motor neuron subtypes. By combining our comprehensive atlas of gastric myenteric neurons with published vagal motor neuron profiles, our work establishes a foundation for systematically dissecting the circuit logic underlying autonomic control of gastric function.

### Sensory response of myenteric neurons in the small intestine

Sensory neurons monitor the state of internal organs. Gastrointestinal distension, intestinal nutrients, bioproduct from gut microbiota and cytokines from the gut immune system have been implicated as sensory cues that activate neurons to regulate gut physiology and animal behavior^3–5,7,56^. Recently advances have primarily focused on extrinsic sensory pathways. Genetically defined vagal sensory neurons form labeled-line circuits for detecting gastric distension and intestinal nutrients^18^. In contrast, while intrinsic primary afferent neurons (IPANs) have been recognized for decades, their capacity to sense and discriminate diverse luminal signals remains poorly understood.

To address this gap, we developed an *ex vivo* calcium imaging preparation capable of monitoring individual enteric neuron responses to multiple luminal stimuli. We discovered that myenteric neurons in the small intestine are activated by nutrients, short-chain fatty acids (enriched in microbial metabolites), irritants, and cytokines. Strikingly, most myenteric neurons exhibited multimodal responsiveness—a coding logic distinct from the taste system but reminiscent of vagal sensory neurons innervating the gut^20,29^. This parallel is biologically plausible, as both enteric and vagal sensory neurons rely on specialized chemosensory cells (e.g., enterochromaffin cells) for luminal nutrient detection. Indeed, we identified functional communication between enterochromaffin cells and enteric neurons. Furthermore, pharmacological inhibition of 5-HT3 and P2X receptors attenuated, but did not abolish, myenteric neuron responses to luminal stimuli, consistent with the receptor expression profile of *Advillin^+^* putative IPANs. The functional significance of *Advillin^+^* neurons in luminal sensing was further underscored by targeted disruption of acetylcholine (ACh) signaling. Genetic deletion of *Chat* (choline acetyltransferase), the key enzyme for ACh synthesis in *Advillin^+^* neurons, significantly attenuated myenteric neuron responses to multiple luminal stimuli. This finding directly implicates cholinergic transmission from *Advillin^+^* neurons as a critical amplifier of intrinsic sensory processing in the enteric nervous system.

Our study extends recent work by Fung et al., who employed similar approaches to profile nutrient responses in enteric neurons of the small intestine^34^. While their work achieved spatial resolution by imaging myenteric and submucosal neurons separately, we focused specifically on decoding calcium dynamics in myenteric circuits. Furthermore, whereas their study combined calcium imaging with post hoc neurochemical marker staining to characterize nutrient-responsive neurons, we broadened the scope of interrogation to include not only more nutrients types but also luminal irritants and immune-derived cytokines. Given the established role of IPANs in mediating gut-immune crosstalk — particularly through bidirectional communication with resident immune cells like ILC2s—our findings provide direct functional evidence for cytokine-triggered activation of myenteric neurons, underscoring the physiological relevance of neuroimmune interactions in gut sensory encoding. Moving forward, integrating multiplexed neurochemical profiling (e.g., multiplex FISH) with iterative *ex vivo* calcium imaging could systematically resolve how distinct luminal signals are discriminatively processed by enteric neural networks.

### Genetic control of enteric IPANs

The ability of the gastrointestinal tract to respond to stimuli ex vivo demonstrates the presence of functional, self-contained neuronal circuits within the enteric nervous system (ENS) capable of initiating sensory-motor reflexes without extrinsic input. Although decades of research have characterized the morphological, neurochemical, and electrophysiological properties of intrinsic primary afferent neurons (IPANs) across mammalian species^57,58^, direct functional manipulation of IPANs within an intact ENS has been needed to definitively establish their role in intrinsic reflex pathways.

In this study, we developed a versatile genetic toolkit to selectively target and control IPANs both *ex vivo* and *in vivo*. Using optogenetic activation, we showed that IPAN stimulation alone is sufficient to elicit region-specific motility changes in the gastrointestinal tract—providing the first direct functional evidence supporting the long-standing model that IPANs can drive motility independently within enteric circuits. Furthermore, genetic ablation of acetylcholine (ACh) synthesis in IPANs impaired gut motility in vivo, underscoring the essential role of these sensory neurons in ENS function. Together, our genetic and functional approaches establish a foundation for mechanistic dissection of defined enteric neuron populations *in vivo*.

## SUPPLEMENTAL FIGURES

**Extended Data Figure 1.**
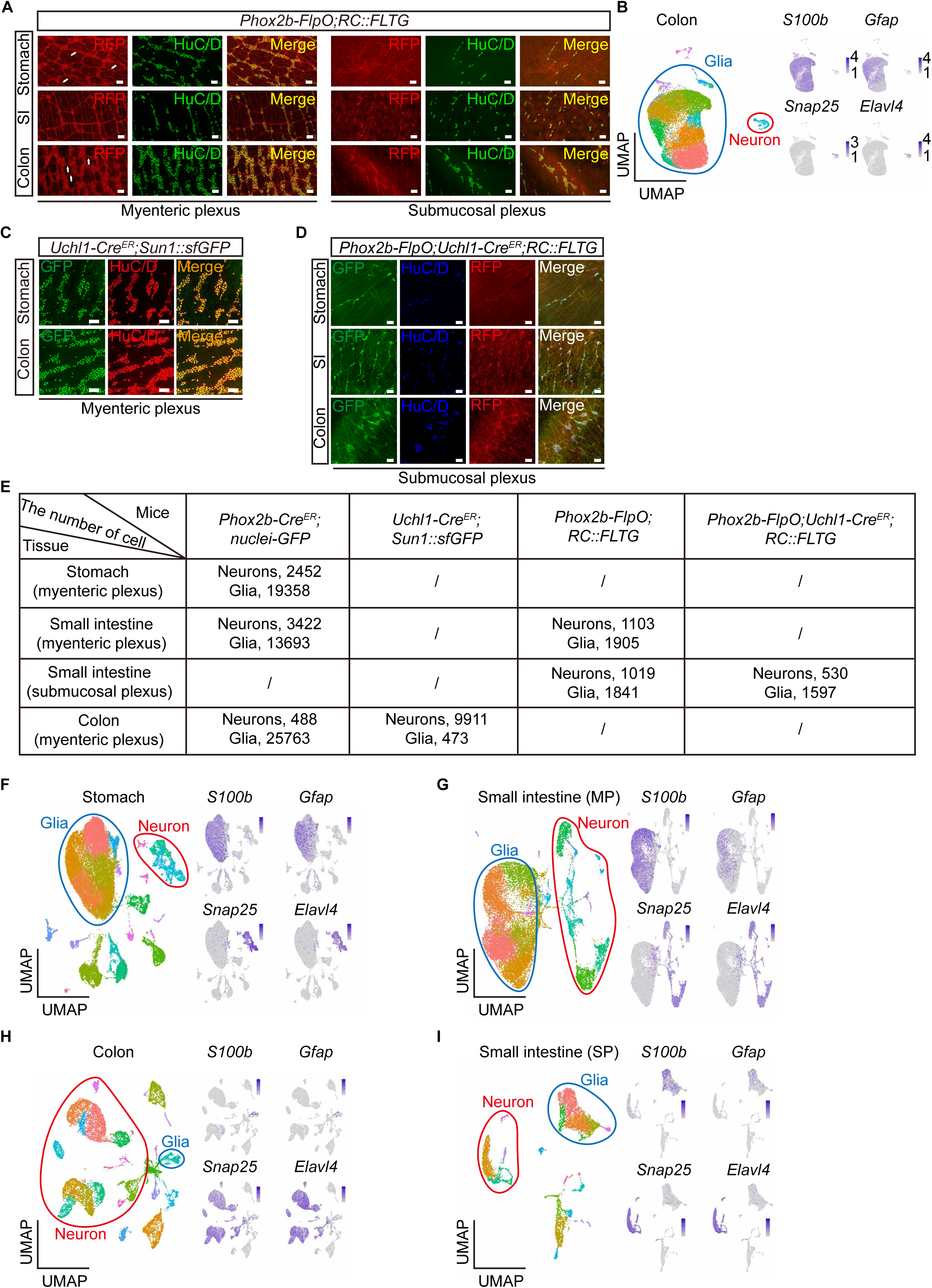
Transgenic tools for segment specific profiling of the mouse ENS, related to Figure 1. **(a)** Representative images for myenteric plexus and submucosal plexus labeled by *Phox2b-FlpO*. HuC/D is used to label enteric neurons. The RC::FLTG is a dual-recombinase reporter. Flp recombinase results in high tdTomato fluorescence, and further exposure to Cre recombinase results in robust eGFP fluorescence. The white arrows indicate inter-ganglionic fiber tracts. Scale bar, 100 μm. **(b)** Uniform manifold approximation and projection (UMAP) results showing neurons and glial cells in the myenteric plexus of colon from *Phox2b-Cre^ER^;Sun1::sfGFP* mice. *Snap25* and *Elavl4* are marker genes for enteric neurons, *S100b* and *Gfap* are marker genes for enteric glia. **(c-d)** Representative images for myenteric or submucosal plexus labeled using indicated reporter lines. HuC/D is used to label enteric neurons. Scale bar, 100 μm. **(e)** Summary of enteric neurons and enteric glia from different segments of the mouse gastrointestinal tract obtained from our scRNA-seq pipeline using different transgenic animal model. **(f-i)** Uniform manifold approximation and projection (UMAP) results showing neurons and glia in the myenteric plexus of the stomach (**f**), small intestine (**g**), colon (**h**), and submucosal plexus of the small intestine (**i**). *Snap25* and *Elavl4* are marker genes for enteric neurons, *S100b* and *Gfap* are marker genes for enteric glia.

**Extended Data Figure 2.**
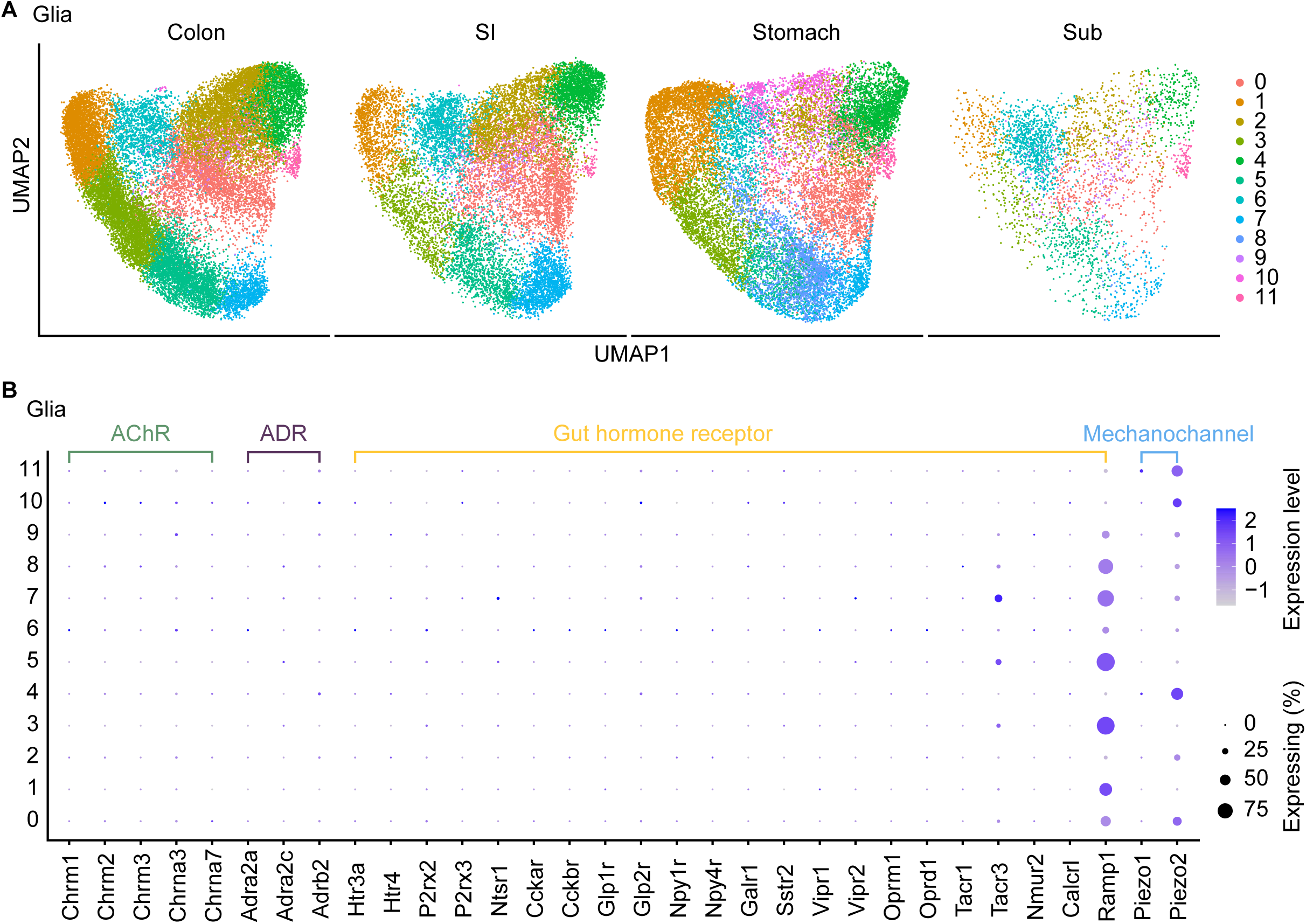
Clustering analysis and molecular profile of enteric glia, related to Figure 1. **(a)** UMAP showing the clustering analysis of enteric glia using scRNA-seq data from myenteric plexus of stomach, small intestine, and colon, as well as submucosal plexus of small intestine, after batch correction using CCAIntegration. **(b)** Differential expressions of receptors and ion channels in enteric glia. Receptors for autonomic neuron transmitters and gut peptides are not detected in enteric glia.

**Extended Data Figure 3.**
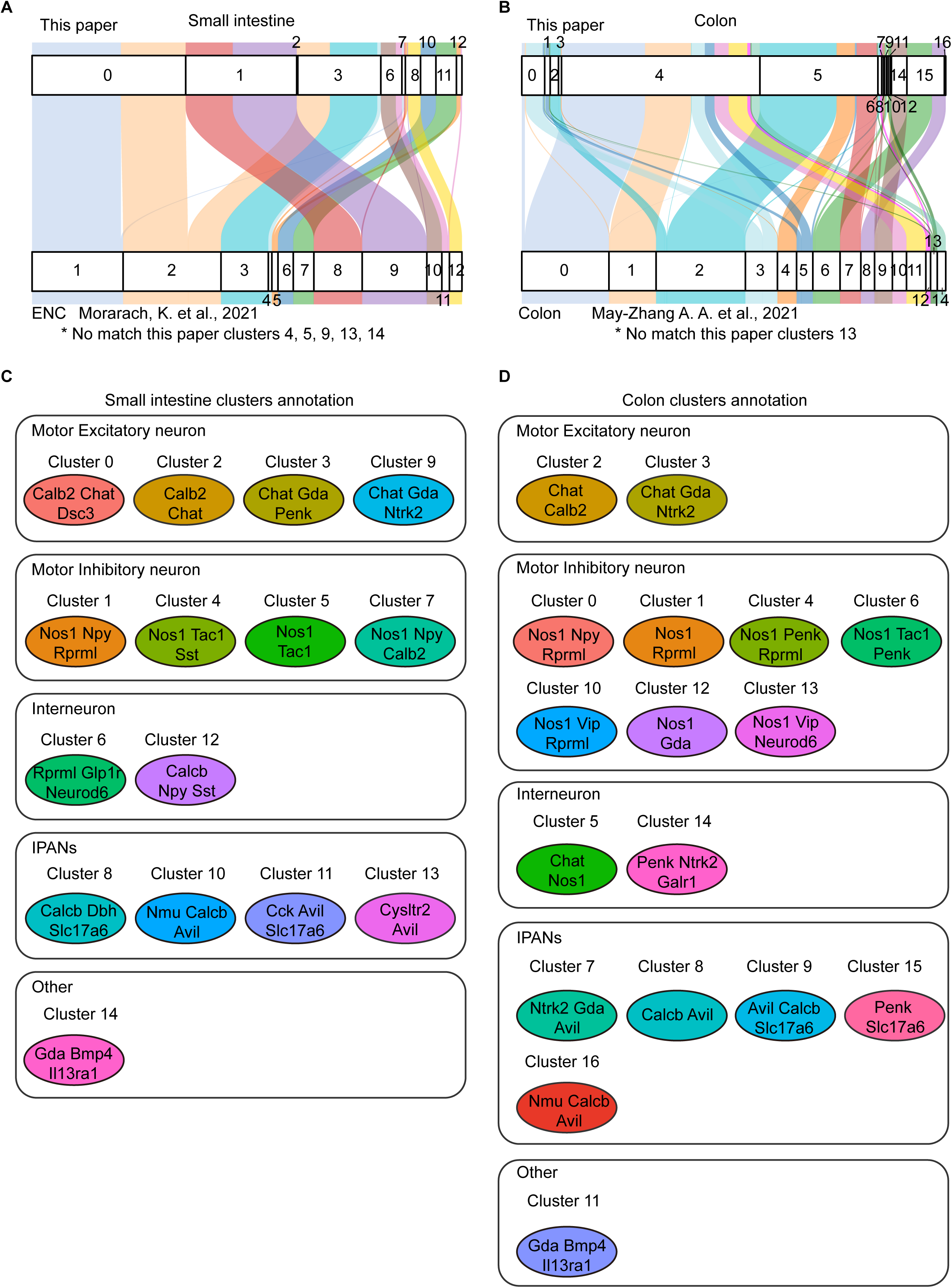
Comparison of ENS scRNA profiling, related to Figure 1. **(a-b)** Label transfer relationship between cell types from our dataset and previously reported dataset. Data for myenteric neurons of the small intestine is from Morarach, K. et al., 2021 and data for myenteric neurons of the colon is from May-Zhang A. A. et al., 2021. **(c-d)** Schematic indicating the proposed functional assignment and selected combinatorial marker genes for each cluster of small intestine and colon in our dataset.

**Extended Data Figure 4.**
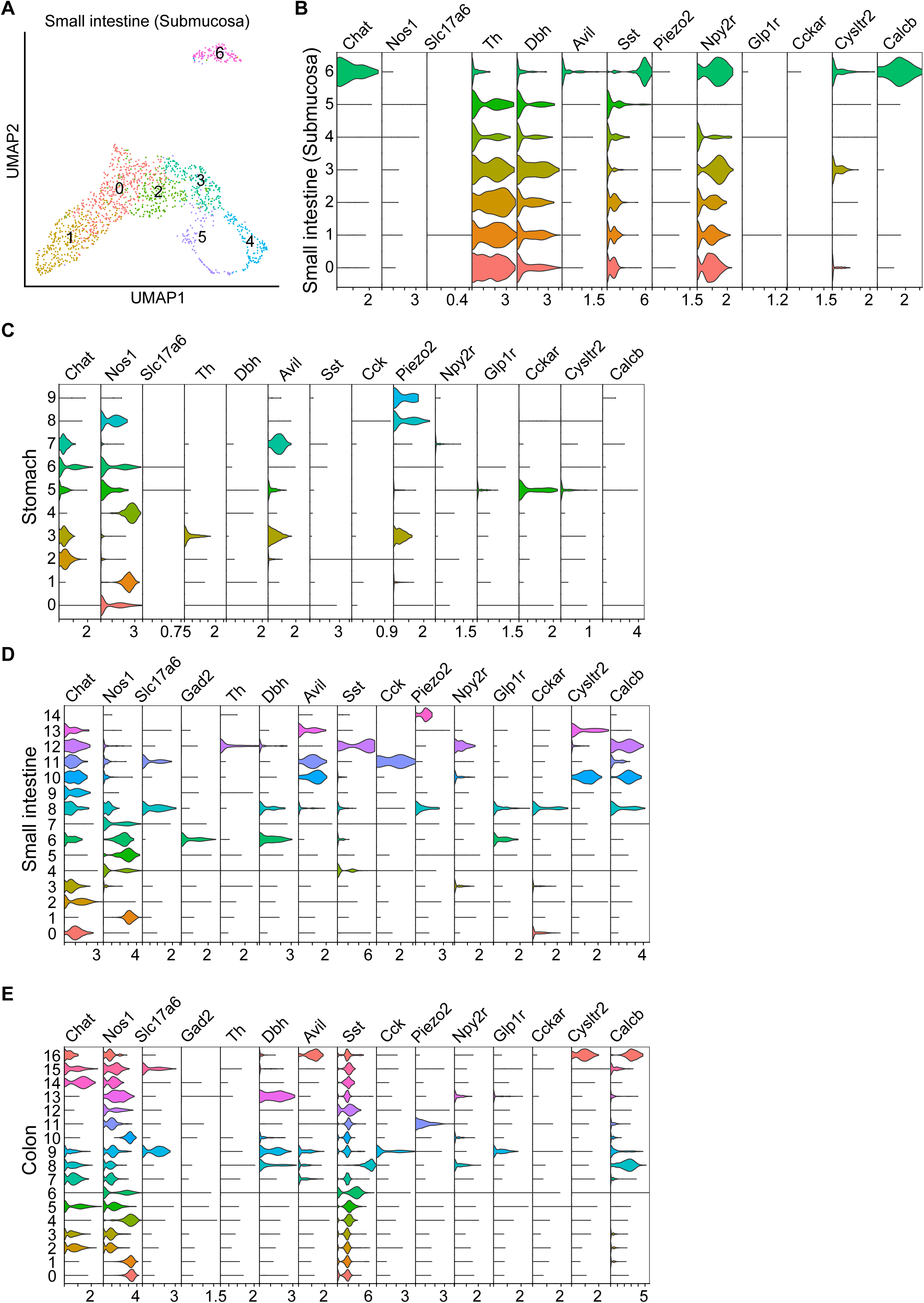
Molecular profiling reveals heterogeneity across various segments of the gastrointestinal tract, related to Figure 1 and Figure 2. **(a)** Re-clustering analysis of submucosal neurons from the small intestine, color-coded to represent 7 distinct clusters. **(b)** VlnPlot showing the expression of selected genes in different clusters of small intestine submucosal neurons. **(c)** VlnPlot showing the expression of selected genes across different clusters of stomach myenteric neurons. **(d)** VlnPlot showing the expression of selected genes across different clusters of small intestine myenteric neurons. **(e)** VlnPlot showing the expression of selected genes in different clusters of colon neurons.

**Extended Data Figure 5.**
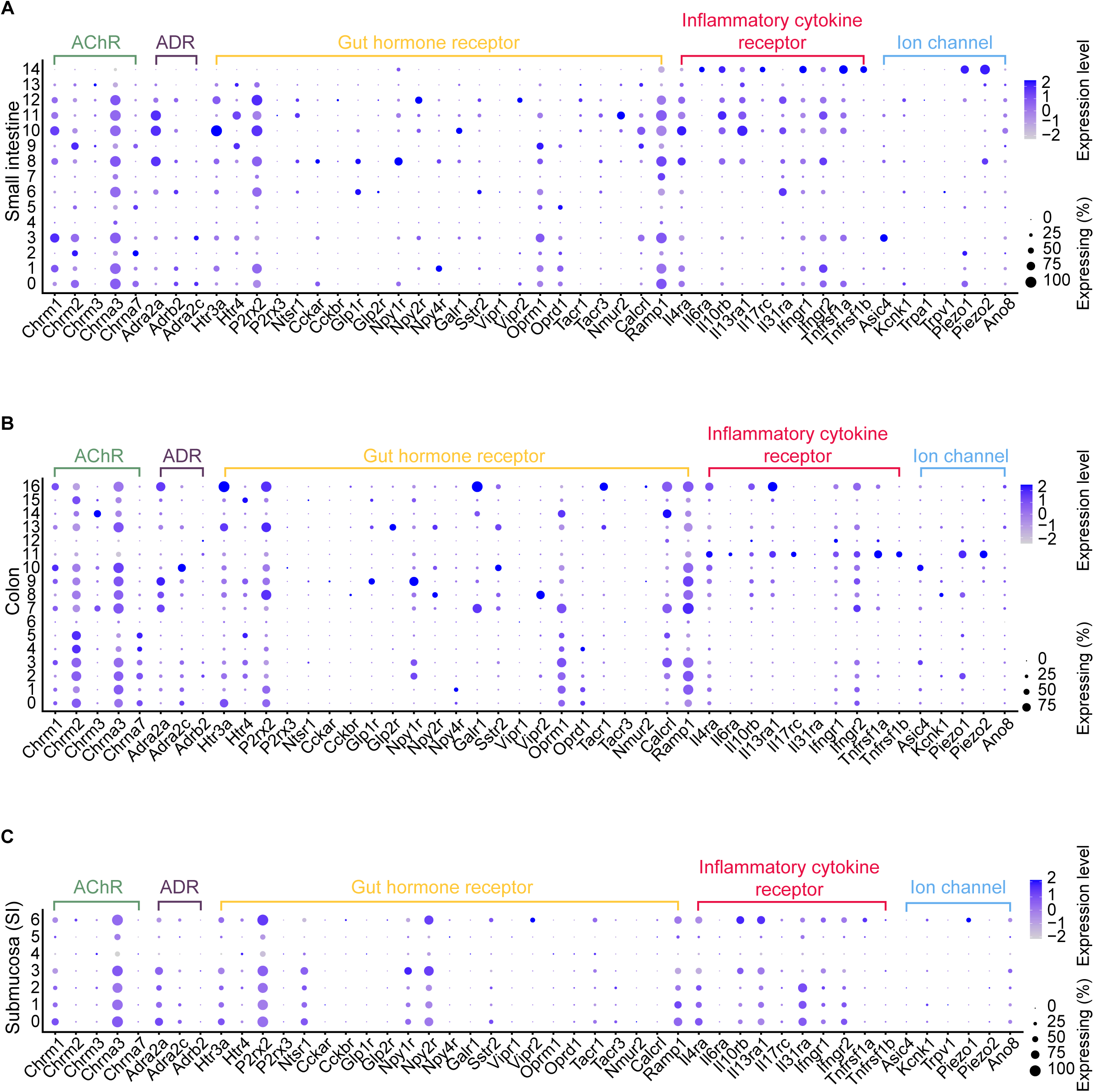
Differential expression of receptors and ion channels in enteric neurons, related to Figure 2. **(a-c)** Dot plots showing differentially expressed genes categorized as acetylcholine receptor (AChRs), adrenergic receptor (ADRs), gut hormone receptors, inflammatory cytokine receptors, and ion channels of enteric neurons in gastrointestinal tract. The color scale represents the gene expression, and dot size represents the percentage of cells.

**Extended Data Figure 6.**
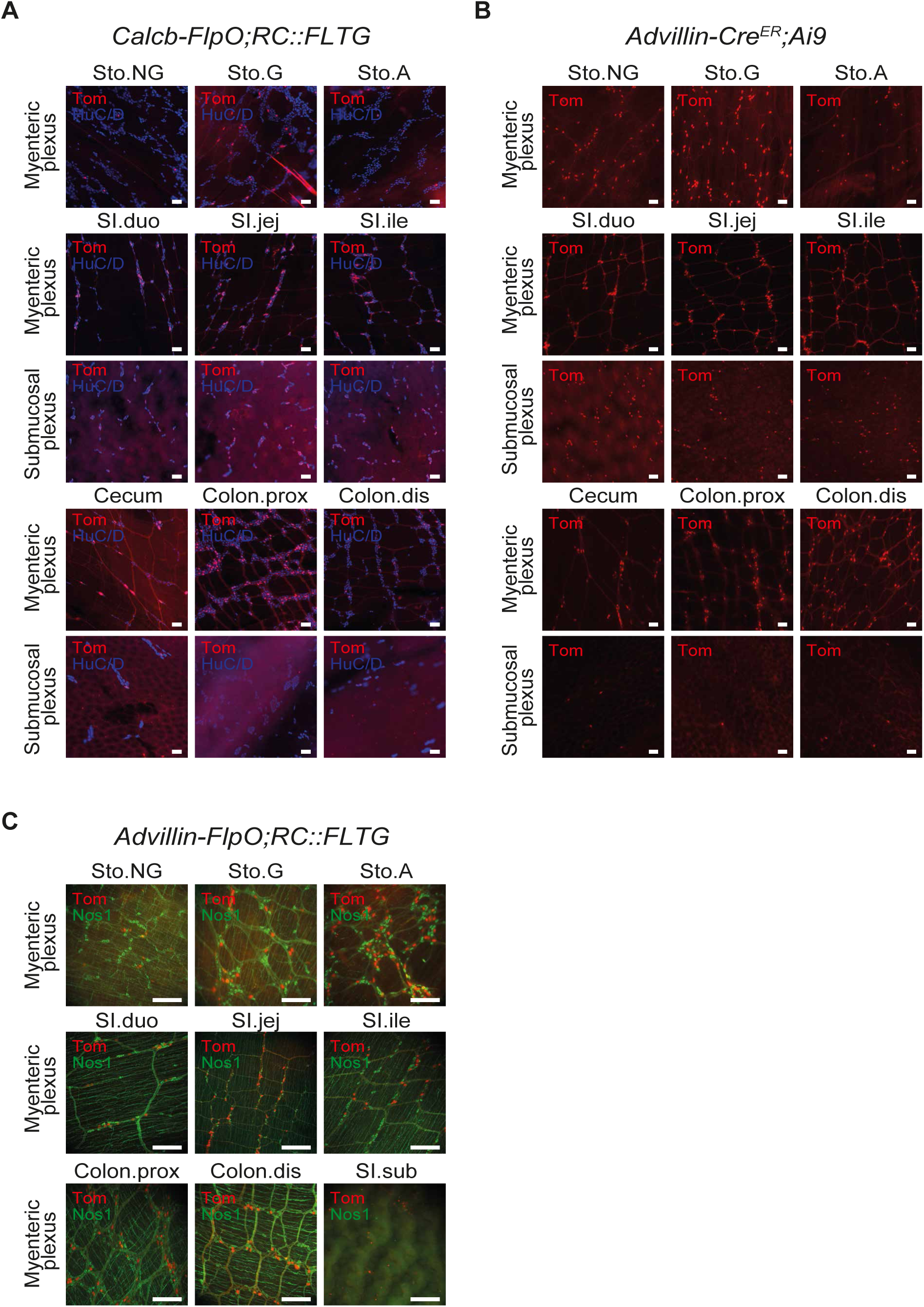
Expression pattern of *Advillin* and *Calcb* reporters, related to Figure 2 and Figure 5 **(a)** Representative images for myenteric plexus and submucosal plexus labeled by *Calcb-FlpO*. HuC/D is used to label enteric neurons. The RC::FLTG is a dual-recombinase reporter. Flp recombinase results in high tdTomato fluorescence, and further exposure to Cre recombinase results in robust eGFP fluorescence. Scale bar, 100 μm. **(b)** Representative images for myenteric plexus and submucosal plexus labeled by *Advillin-Cre^ER^*. Ai9 reporter expressing Cre dependent tdTomato is used to visualize *Advillin-Cre^ER^* cells. Scale bar, 100 μm. **(c)** Representative images for myenteric plexus labeled by *Advillin-FlpO*. The RC::FLTG is used to visualize *Advillin-FlpO* cells Scale bar, 100 μm.

**Extended Data Figure 7.**
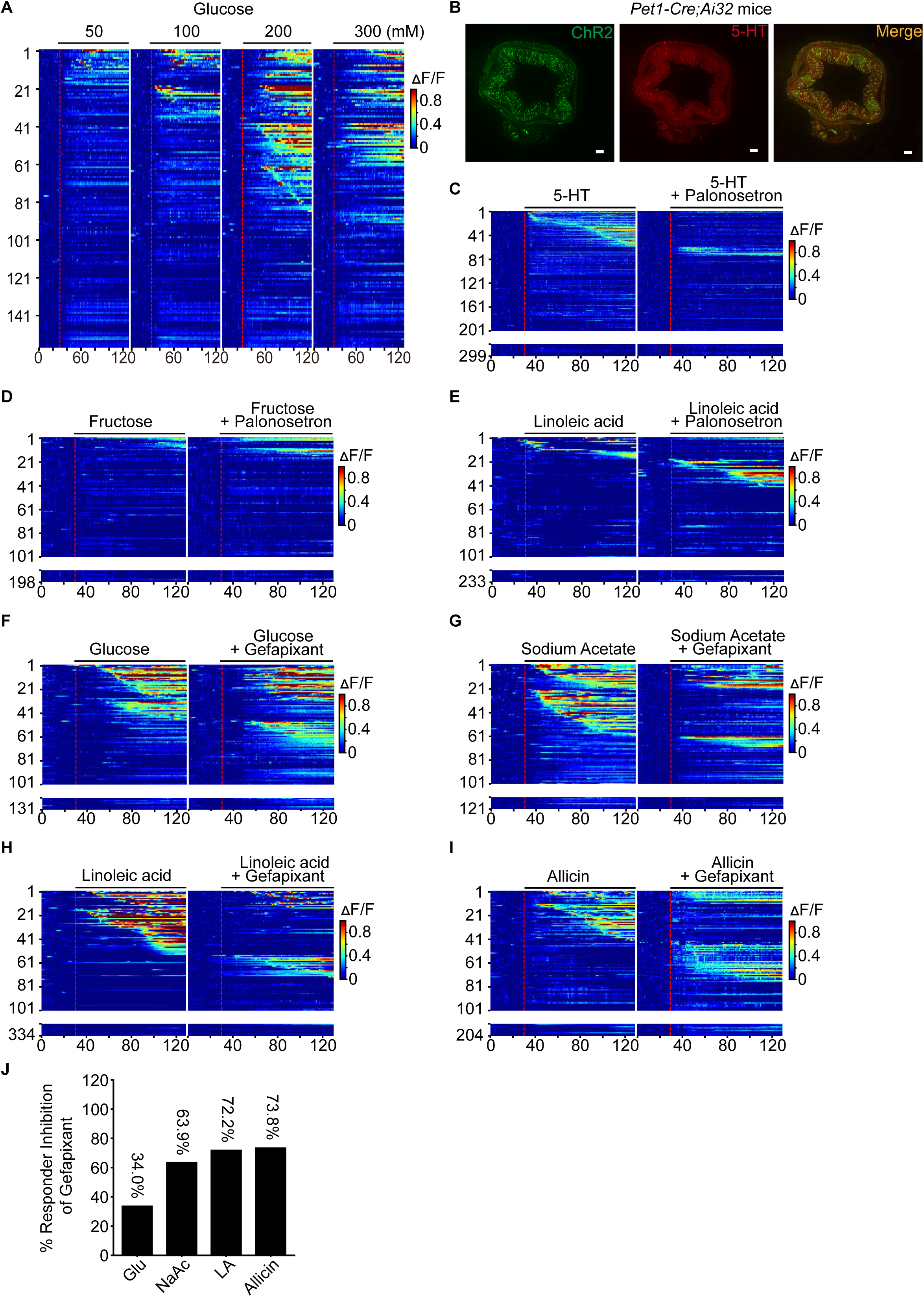
Response of enteric neurons to luminal stimuli, related to Figure 3 and Figure 4. **(a)** Calcium responses of enteric neurons to different concentrations of glucose. The red dashed line in the heatmap indicates the administration of the stimuli. Each row represents the response from the same neuron. **(b)** Cross sections showing ChR2 expression in 5-HTL enterochromaffin cells of *Pet1;Ai32* mice. Scale bar, 200 μm. **(c-e)** Calcium responses of enteric neurons to 100 μM 5-HT (**c**), 125 mM fructose (**d**), and 100mM linoleic acid (**e**) before and after 1 μM Palonosetron (an antagonist of 5-HT_3_R) treatment. The red dashed line indicates the administration of the stimuli. Palonosetron were incubated for the luminal side 5 min before recording and were maintained throughout the recording session. **(f-i)** Calcium responses of enteric neurons to 125 mM glucose (**f**), and 50 mM sodium acetate (**g**), 100 mM linoleic acid (**h**), and 0.5% allicin (**i**) before and after 1 μM Gefapixant (an antagonist of P2X3R) treatment. The red dashed line indicates the administration of the stimuli. Gefapixant were incubated for the luminal side 5 min before recording and were maintained throughout the recording session. **(j)** Quantification for proportion of responders inhibited by Gefapixant.

**Extended Data Figure 8.**
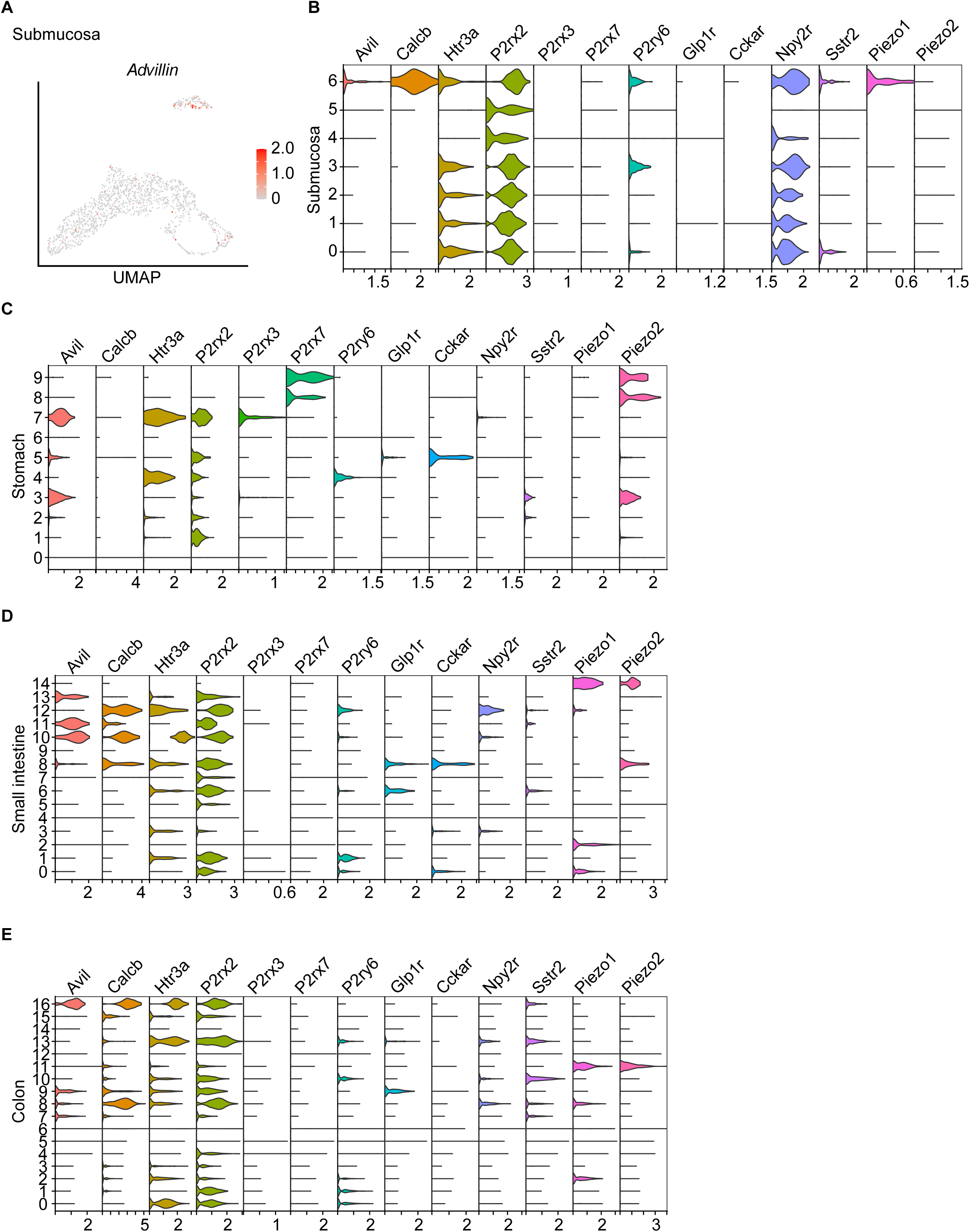
Expression pattern of gut peptide receptors and mechanosensitive ion channels in IPANs, related to Figure 5. **(a)** Feature plot showing the expression of *Advillin* in submucosa plexus of the small intestine. **(b-e)** Volin plot showing the expression of indicated receptor gene in the submucosal plexus of small intestine (**b**), as well as myenteric plexus of stomach (**c**), small intestine (**d**) and colon (**e**). Note the expression of 5-HT receptors, ATP receptors, gut peptide receptors and Piezo2 in *Advillin^+^* (Avil) myenteric neurons.

**Extended Data Figure 9.**
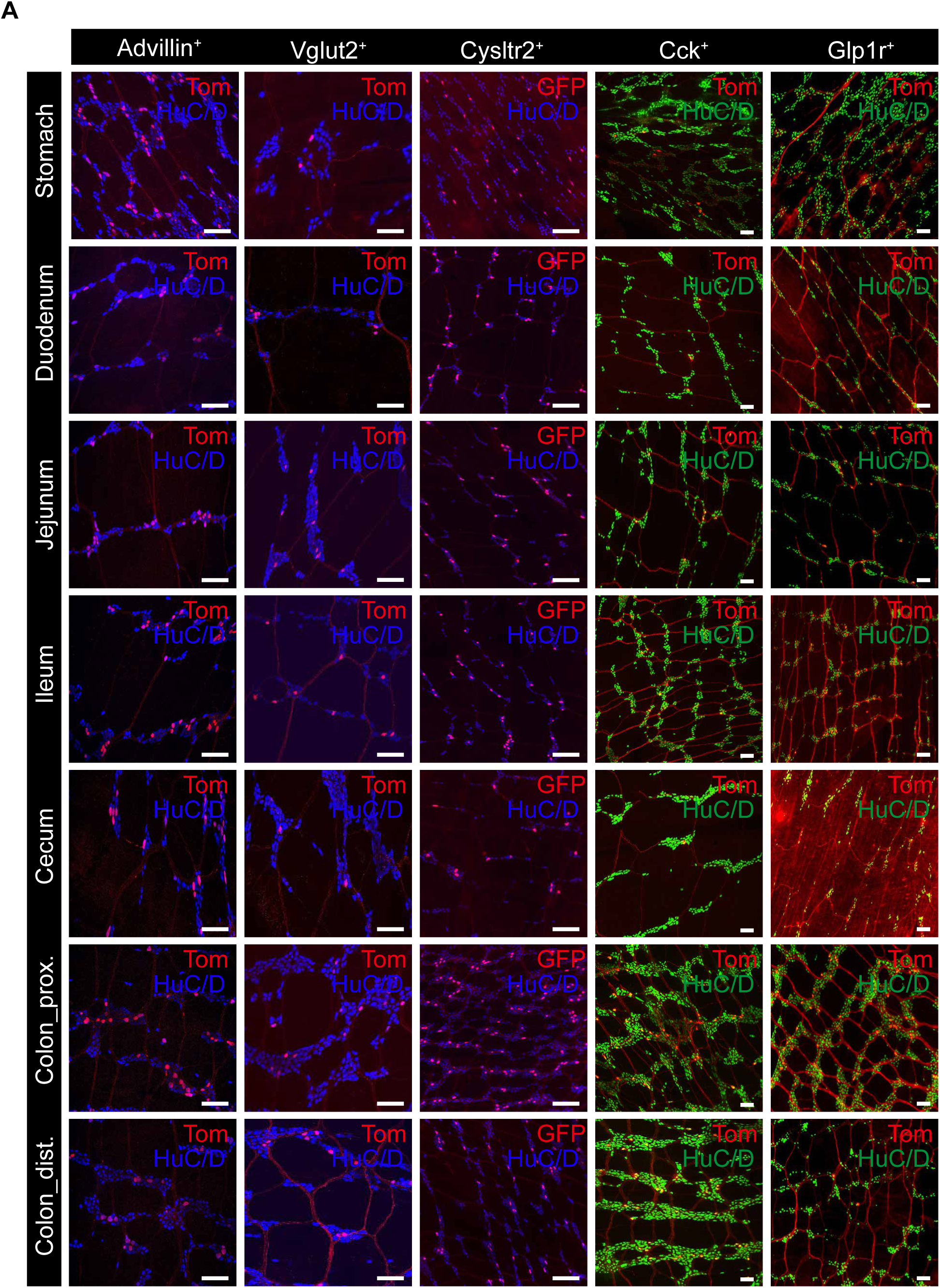
Visualization of different subsets of *Advillin^+^* myenteric IPANs by intersectional genetics, related to Figure 5. **(a)** Images showing the distribution of myenteric neurons labeled by indicated genetic markers in the gastrointestinal tract. Genotypes of mice used for labeling each genetic markers: Advillin (*Avil-FlpO^+/-^, RC::FLTG^+/-^*), Vglut2 (*Vglut2-Cre^+/-^*, *Ai9^+/-^*), Cysltr2 (*Phox2b-FlpO^+/-^*, *Cystrl2-Cre^+/-^*, *RC::FLTG^+/-^*), Cck (*Vglut2-FlpO^+,/-^*, *Cck-Cre^+/-^*, *Ai65^+/-^*), Glp1r (*Vglut2-FlpO^+,/-^*, *Glp1r-Cre^+/-^*, *Ai65^+/-^*). The RC::FLTG is a dual-recombinase reporter. FlpO recombinase results in high tdTomato fluorescence, and further exposure to Cre recombinase results in robust eGFP fluorescence. Ai65 is a dual-recombinase reporter in which exposure to FlpO and Cre result in the expression of tdTomato. Scale bar: 100 μm

**Extended Data Figure 10.**
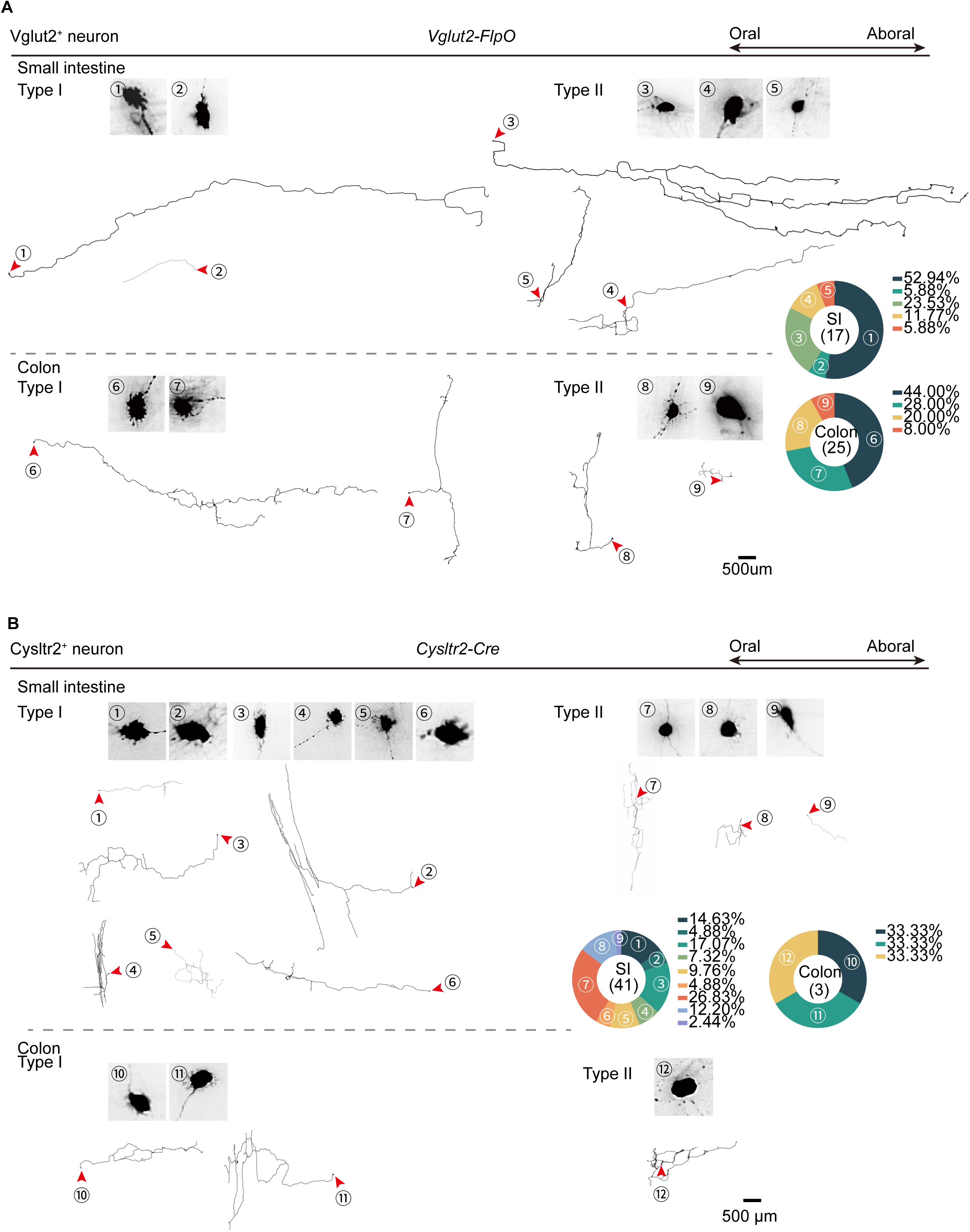
Morphological diversity of *Vglut2^+^*, *Cysltr2^+^* myenteric neurons, related to Figure 5. **(a)** Reconstructed single neuron morphology of *Vglut2^+^* myenteric neurons from the small intestine and colon. A total of 42 neurons with 9 morphological types are analyzed by sparse labeling. The distribution of each morphological type shown is quantified in small intestine and colon separately and shown in the pie chart. Scale bar, 500 μm. **(b)** Reconstructed single neuron morphology of *Cysltr2^+^* myenteric neurons from the small intestine and colon. A total of 44 neurons with 12 morphological types are analyzed by sparse labeling. The distribution of each morphological type shown is quantified in small intestine and colon separately and shown in the pie chart. Scale bar, 500 µm

**Extended Data Figure 11.**
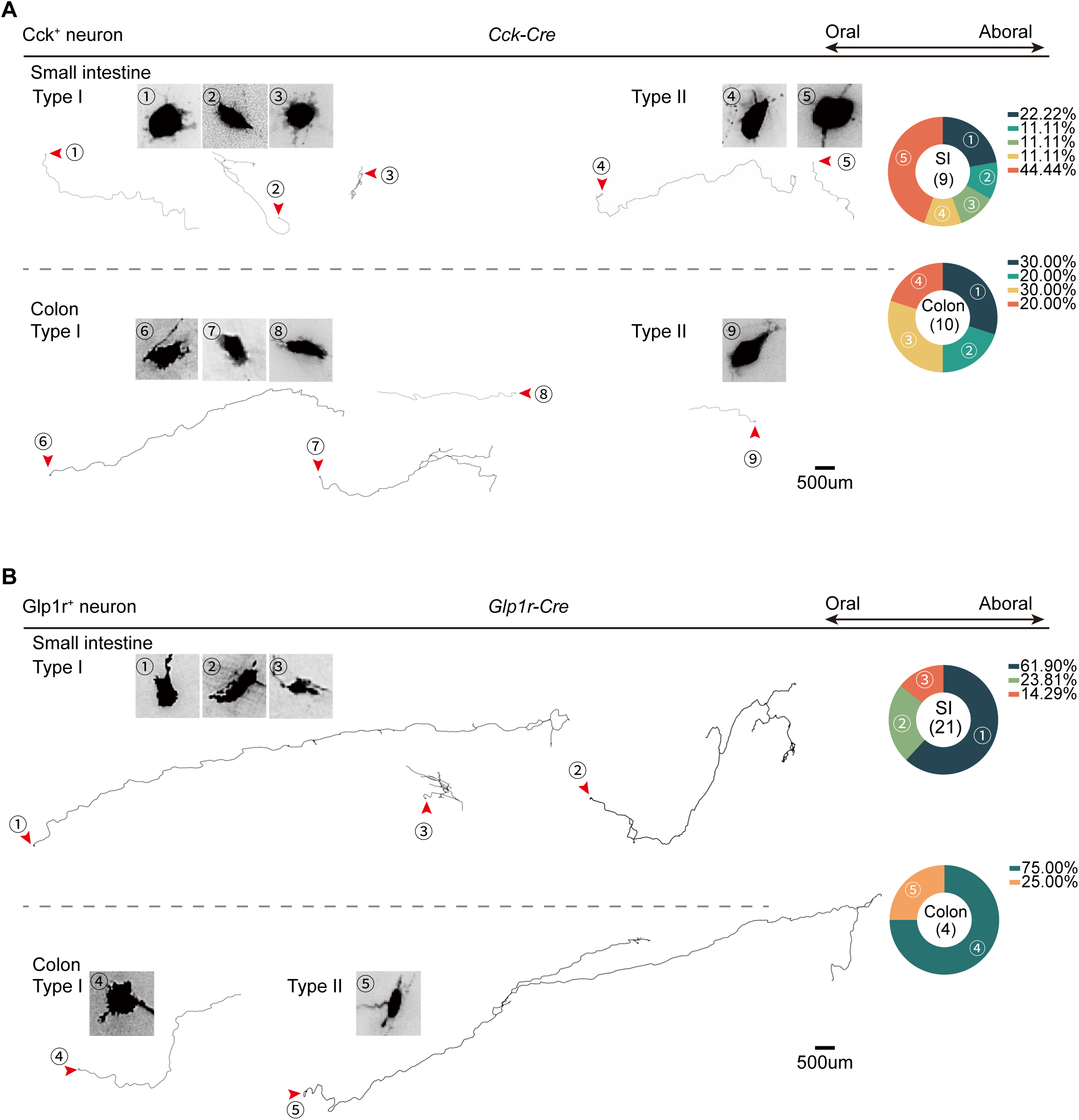
Morphological diversity of *Cck^+^* and *Glp1r^+^* myenteric neurons, related to Figure 5. **(a)** Reconstructed single neuron morphology of *Cck^+^*myenteric neurons from the small intestine and colon. A total of 19 neurons with 9 morphological types are analyzed by sparse labeling. The distribution of each morphological type shown is quantified in small intestine and colon separately and shown in the pie chart. Scale bar, 500 μm. **(b)** Reconstructed single neuron morphology of *Glp1r^+^* myenteric neurons from the small intestine and colon. A total of 25 neurons with 5 morphological types are analyzed by sparse labeling. The distribution of each morphological type shown is quantified in small intestine and colon separately and shown in the pie chart. Scale bar, 500 μm.

**Extended Data Figure 12.**
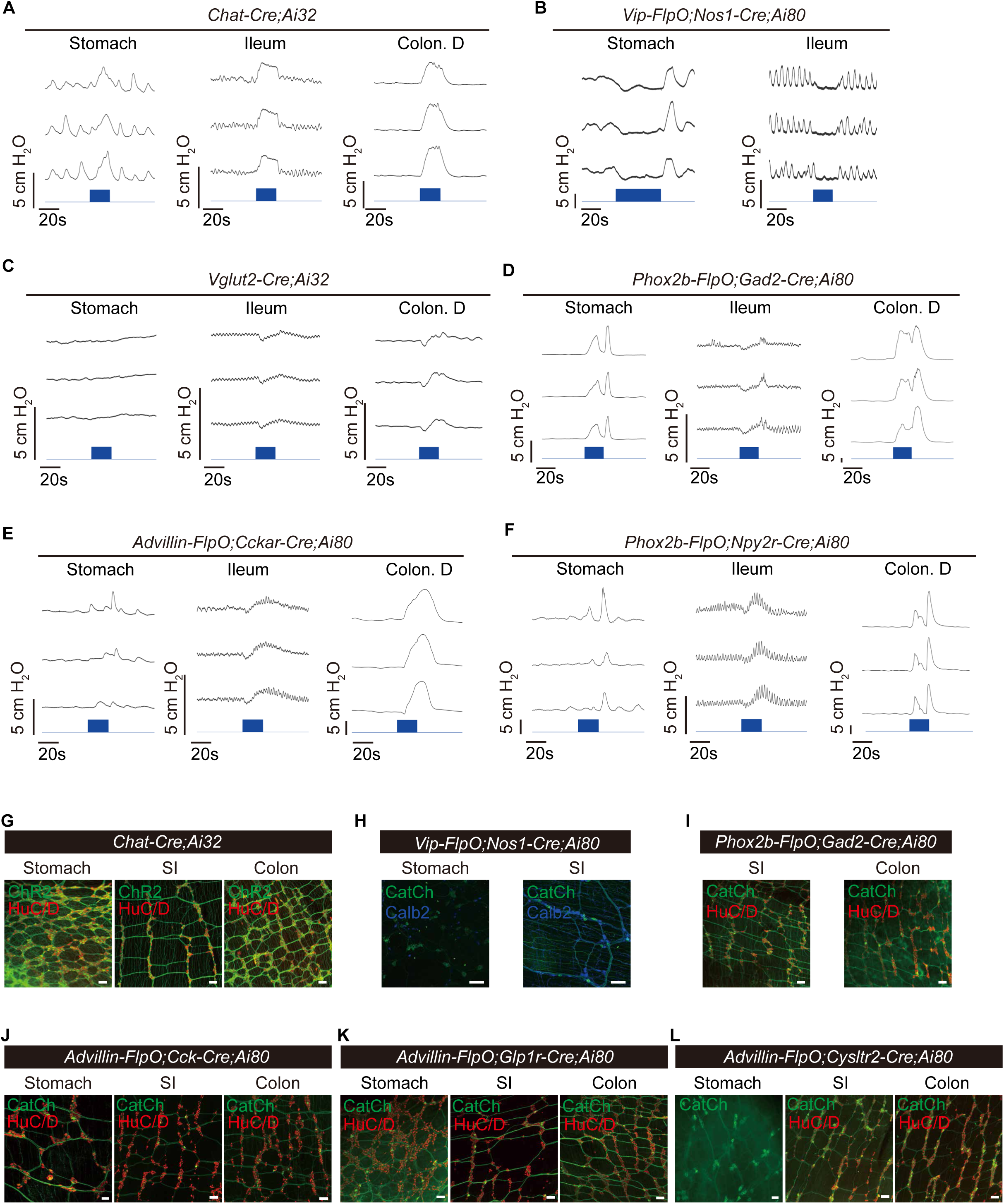
Optogenetic control of gut motility by enteric neurons, related to Figure 7. **(a-f)** Representative traces showing that optogenetic induced gut motility change. Gut motility change is reflected by alteration in luminal pressure (centimeter H_2_O pressure) measured by the pressure sensor. Blue light bar in the bottom indicates laser stimulation. **(g-l)** Immunofluorescence images showing the expression of ChR2 or CatCh in myenteric neurons by indicated genetic reporters in the gastrointestinal tract. HuC/D or Calb2 is used to label enteric neurons. Scale bar, 100 μm. Ai80 (RCFL-CatCh) is a dual-recombinase reporter mice which drives CatCh-EYFP expression in FlpO and Cre double positive cells.

